# Phenotypic and genomic changes during *Turnip mosaic virus* adaptation to *Arabidopsis thaliana* mutants lacking epigenetic regulatory factors

**DOI:** 10.1101/2023.05.17.541084

**Authors:** Silvia Ambrós, María J. Olmo-Uceda, Régis L. Corrêa, Santiago F. Elena

## Abstract

In this study we investigated how RNA viral populations evolve, interact and adapt to epigenetically-controlled plant defense mechanisms. We have evolved five independent lineages of turnip mosaic virus (TuMV) in a set of *Arabidopsis thaliana* genotypes carrying mutations that influence important elements of two main epigenetic pathways. All evolved lineages showed adaptation to the lack of epigenetically-regulated responses through significant increases in infectivity, virulence and viral load although the magnitude of the improvements strongly depended on the plant genotype. In early passages, these traits evolved more rapidly, but the rate of evolution flattened out in later ones. Viral load was positively correlated with different measures of virulence, though the strength of the associations changed from the ancestral to the evolved viruses. High-throughput sequencing was used to evaluate the viral diversity of each lineage, as well as characterizing the nature of fixed mutations, evolutionary convergences and potential targets of TuMV adaptation. Within each lineage, we observed a net increase in genome-wide genetic diversity, with some instances where nonsynonymous alleles experienced a transient rise in abundance before being displaced by the ancestral allele. Viral VPg protein has been shown as a key player in the adaptation process, even though no obvious association between fixed alleles and host genotype was found.

**Layman Summary:** Epigenetic factors influence the expression of defense genes in plants, allowing for phenotypic rapid responses to infections by pathogens. The role of epigenetics in shaping the coevolution between host and pathogens has received very little attention. Here, we explored how RNA viruses interact and adapt to plant defense mechanisms that are controlled by epigenetic factors. We conducted evolution experiments on turnip mosaic virus using *Arabidopsis thaliana* genotypes with mutations that affect epigenetic pathways. We found that all evolved viral lineages adapted to the alteration of epigenetically-regulated responses by becoming more infectious, virulent, and having a higher viral load. The improvements varied depending on the plant genotype. The study also found that viral load was positively correlated with virulence, but the associations changed from the original to the evolved viruses. We used high-throughput sequencing to evaluate viral diversity and found an increase in each evolving lineage. We found that virus adaptation primarily targeted viral VPg, despite no obvious association between fixed alleles and host genotype being found.

**Teaser Text:** Discover how RNA viruses adapt and evolve to plant defense mechanisms controlled by epigenetic factors. This research found that epigenetic regulation of defense genes modulates viral evolution. Viral lineages became more infectious, virulent, and had a higher viral load. Find out more about the correlation between viral load and virulence, viral diversity, and the primary virus genomic target of adaptation.

## Introduction

Plants and viruses interact, triggering defense and counter-defense mechanisms that frequently lead to coevolution and reciprocal adaptation. A variety of plant immune processes participate in antiviral defense (Soosaar et al. 2005; Zhou and Zhang 2020). These antiviral factors can be broadly divided into two groups: (*i*) basal, which are preexisting and restrict within-cell propagation and cell-to-cell spread, and (*ii*) inducible, which are triggered following infection and inhibit systemic movement and replication. Inducible mechanisms involve genes whose expression results in a broad-scale change in plant physiology via a variety of signaling pathways, as opposed to basal mechanisms, which correspond to alleles of cell proteins whose interaction with viral factors is altered (Carr et al. 2010). These changes include local cell apoptosis (Loebenstein 2009), upregulation of nonspecific responses against pathogens throughout the entire plant (systemic acquired and induced resistances) (Kachroo et al. 2006; Carr et al. 2010) and the activation of the RNA-silencing-based resistance, that seems to play a role both in basal and inducible mechanisms (Voinnet 2001; Carr et al. 2010; López-Gomollon and Baulcombe 2022). RNA-based immunity is triggered by the recognition and degradation of double-stranded RNAs (dsRNA). Viral dsRNAs produced during replication are degraded by Dicer-like (DCL) proteins into small RNAs (sRNA) that are loaded into Argonaute (AGO) proteins and used as guide to repress complementary RNAs (Voinnet 2001). The silencing response can be enhanced by RNA-dependent RNA polymerases (RDR), which produce extra dsRNAs from targets (Borges and Martienssen 2015).

The RNA-mediated mechanisms employed by plants to defend against viruses are a component of a larger and evolutionarily conserved system that governs gene expression and manages transposable elements (TE) via the addition of epigenetic modifications to DNA or histones (Hung and Slotkin 2021). DNA methylation can be observed in all sequence contexts in eukaryotes (CG, CHG and CHH; with H being any nucleotide except G). RNA-directed DNA methylation (RdDM) is a well-studied mechanism that uses sRNAs produced from the borders of TEs to restrict their expression, influencing the expression of neighboring genes (Liu et al. 2022). RdDM plays an important role in determining heterochromatin boundaries, coordinating the actions of components such as METHYLTRANSFERASE DOMAINS REARRANGED METHYLASE 2 (DRM2), RNA polymerase IV (POLIV), and RNA polymerase V (POLV) (Böhmdorfer et al. 2016). Alternative non-canonical pathways (*e.g*., POLII-derived mRNAs, RDR6-derived dsRNAs, or DCL-independent mechanisms) feed into the canonical RdDM pathway (Cuerda-Gil and Slotkin 2016). Gene expression and TE mobilization can also be controlled via sRNA-independent mechanisms in which epigenetic marks are copied during replication by maintenance DNA methyltransferases such as the plant-specific CHROMOMETHYLASE 3 and require chromatin remodelers such as DECREASED DNA METHYLATION 1 (DDM1) (Bond and Baulcombe 2014). The introduction of changes in histones is commonly related with the targets of DNA methylation marks. Histone H3 trimethylation of lysine 9 (H3K9m3), for example, has been linked to TE repression, whereas H3K4m3 has been linked to gene expression stimulation. DNA methylation and histone modification are both reversible markers. The proteins REPRESSOR OF SILENCING 1 (ROS1), INCREASE IN BONSAI METHYLATION 1 (IBM1), and JUMONJI14 (JMJ14), for example, are involved in the demethylation of DNA, H3K9, and H3K4 respectively (Gong et al. 2002; Saze et al. 2008; Lu et al. 2010). The chromatin environment within or surrounding TEs and genes is altered under stressful situations or in epigenetically-deficient mutants, influencing their expression (Lloyd and Lister 2021).

The link between the loss of DNA methylation factors and resistance or susceptibility to bacteria and fungi has been stablished (*e.g*., López et al. 2011; Dowen et al. 2012; Luna et al. 2012; Yu et al. 2013; Le et al. 2014; López Sánchez et al. 2016). However, the role of RNA virus infection in DNA methylation responses received less attention. Using turnip crinkle virus (Diezma-Navas et al. 2019), two tobamoviruses (Leone et al. 2020), and turnip mosaic virus (TuMV) (Corrêa et al. 2020), it has been shown that *Arabidopsis thaliana* mutants for various RdDM factors, chromatin remodelers and histone modifiers were more or less susceptible to infections depending on the presence of repressive marks. Furthermore, evolution experiments in mutants for innate immunity pathways or basal and inducible defense pathways have shown that viral populations quickly respond to the novel selective pressures imposed by mutations in the RdRM genes (Mongelli et al. 2022; Navarro et al. 2022), suggesting a direct role of these genes in the infection cycle of RNA viruses.

In this work, we leveraged the knowledge on *A. thaliana* epigenetic regulation mechanisms, the intrinsic variability and evolvability of the TuMV mutant swarm, and the great power of high-throughput RNA-Seq to study the impact of epigenetic pathways on viral diversity and evolution. Our results with infections in plants defective for several epigenetic pathways show that the host genotype affects viral genetic diversity independent of the pathway being disrupted, and it is followed by an increase in virulence along evolutionary passages.

## Methods

### Plants, viruses, and growth conditions

A collection of 20 different *A. thaliana* (L.) HEYNH mutants of the Col-0 accession have been used (Table 1). This collection covers a number of single, double and even triple mutants affecting different genes involved in canonical and non-canonical RdDM, sRNA-independent DNA methylation and histone modification. In all experiments, plants were maintained in a BSL2 climatic chamber under a photoperiod of 8 h light (LED tubes at PAR 90 - 100 μmol m^−2^ s^−1^) at 24 °C and 16 h dark at 20 °C and 40% relative humidity.

**Table 1.** Different *A. thaliana* genotypes used in this study. Highlighted in gray are those used for the evolution experiments

| Genotype (allele) | Gene name | Affected pathway and expected phenotype | Reference |
| --- | --- | --- | --- |
| Wild-type (WT) |  |  |  |
| *ago4-2* | *AGO4* | Canonical and DCL-independent RdDM initiation.<br>Reduced non-CpG methylation and gene activation in RdDM targets. | Zilberman et al. (2003) |
| *atx1-2* | *ATX1* | H3K4 histone methyltransferase.<br>Repression of ATX1 targets, including pathogen- and disease-resistance encoding genes | Alvarez-Venegas et al. (2003) |
| *cmt3-11t* | *CMT3* | Maintenance CHG methylation.<br>CHG siRNA-independent DNA methylation lost. | Lindroth et al. (2001) |
| *dcl2-1 dcl3-1 dcl4-2t* | *DCL2, DCL3, DCL4* | siRNAs production.<br>All DCL-dependent RdDM and RNAi pathways compromised. | Bouché et al. (2006) |
| *ddm1-2* | *DDM1* | Chromatin remodeling and stability.<br>All siRNA-independent methylation is compromised in repeated sequences. | Jeddeloh et al. (1999) |
| *ddm1-2 poliv* | *DDM1, NRPD1a* | Chromatin remodeling and stability and canonical RdDM.<br>Only POLII initiation in TE induced situation. | McCue et al. (2012) |
| *ddm1-2 rdr6-15* | *DDM1, RDR6* | Chromatin remodeling and stability and amplification of siRNAs.<br>POLIV-RDR2 pathways working in TE induced situation. | McCue et al. (2012) |
| *ddm1-2 poliv rdr6-15* | *DDM1, NRPD1a, RDR6* | Chromatin remodeling and stability, canonical RdDM and siRNA amplification<br>POLIV and RDR6-dependent RdDM pathways compromised. | Nuthikattu et al. (2013) |
| *drm1-2 drm2-2* | *DRM1, DRM2* | Canonical and non-canonical RdDM.<br>siRNA-dependent CHH lost. | Cao and Jacobsen (2002) |
| *drm1-2 drm2-1 cmt3-7* | *DRM1, DRM2, CMT3* | Canonical and non-canonical RdDM.<br>siRNA-independent and -dependent CHG/CHH lost. | Chan et al. (2006) |
| *hda6-6* | *HDA6* | Histone deacetylase.<br>Increased histone acetylation in HDA6 targets (genes and repeats). | Earley et al. (2006) |
| *ibm1-4* | *IBM1* | H3K9 histone demethylase.<br>H3K9me2 and CHG DNA hypermethylation in a large number of genes. | Saze et al. (2008) |
| *jmj14-1* | *JMJ14* | H3K4 histone demethylase.<br>Increased H3K4me3 and gene activation of JMJ14 targets. | Searle et al. (2010) |
| *poliv (nrpd1a-3)* | *NRPD1a* | Canonical and NERD<sup>1</sup> RdDM.<br>Loss of 24 nt siRNAs associated with *de novo* methylation of repeats. | Herr et al. (2005) |
| *polv (SALK_017795)* | *NRPE1* | Canonical, NERD and RDR6 RdDM.<br>Derepression of TEs mostly from intergenic regions. | Pontier et al. (2005) |
| *rdr1-1 rdr2-1 rdr6-15* | *RDR1, RDR2, RDR6* | RNA silencing, tasiRNA, Canonical RdDM and RDR6-RdDM.<br>Most RNA silencing and RdDM pathways compromised. | Garcia-Ruiz et al. (2010) |
| *rdr2-1* | *RDR2* | Canonical RdDM.<br>Loss of 24 nt siRNA amplification associated with *de novo* methylation of repeats. | Xie et al. (2004) |
| *rdr6-15* | *RDR6* | RNA silencing, tasiRNA and RDR6-RdDM.<br>Unable to amplify sRNAs. POLIV-RDR2 pathway still working. | Mourrain et al. (2000) |
| *ros1-4* | *ROS1* | DNA demethylase. Spread of repressive marks from TEs to nearby genes. | Gong et al. (2002) |

<sup>1</sup>NERD: Needed for RDR2-independent DNA methylation. Binds to H3K4 and triggers siRNA-dependent DNA methylation (Pontier et al. 2012).

TuMV infectious sap was obtained from TuMV-infected *Nicotiana benthamiana* DOMIN plants inoculated with the plasmid p35STunos containing a cDNA of the TuMV YC5 strain genome (GenBank AF30055.2) under the control of the 35S promoter and the *nos* terminator (Chen et al. 2003) as described in Corrêa et al. (2020). After plants showed symptoms of infection, they were pooled and frozen with liquid N_2_. Frozen plant tissue was homogenized using a Mixer Mill MM400 (Retsch GmbH, Haan, Germany). For inoculations, 0.1 g were diluted in 1 ml inoculation buffer (50 mM phosphate buffer pH 7.0, 3% PEG6000, 10% Carborundum) and 5 μL rubbed onto two leaves per plant. Plants were all inoculated at growth stage 3.5 (Boyes et al. 2001). This synchronization ensures that all hosts were at the same phenological stage when inoculated.

### Phenotyping infection

Five different disease-related traits were measured for each plant (20 for preliminary experiments and 10 for evolution experiments) during 14 days post-inoculation (dpi) (17 dpi in case of the second block of preliminary experiments): (*i*) Mean infectivity, as the average frequency of symptomatic plants out of the number of inoculated plants at 14 dpi. (*ii*) Area under the disease progress stairs (*AUDPS*) summarizes in a single figure the temporal dynamics at which symptoms first appear (Simko and Piepho 2012); *AUDPS* ∈ [0, *T*], where *T* is the total number of days in the assay. (*iii*) Average severity of symptoms at 14 dpi, evaluated in a discrete scale ranging from zero for non-infection or asymptomatic infections to five for plant showing generalized necrosis and wilting (Butković et al. 2021). (*iv*) Area under the symptoms intensity progression stair (*AUSIPS*) summarizes both the severity of symptoms and their temporal dynamics; *AUSIPS* ∈ [0, 5*T*]. And (*v*) viral load, measured by absolute reverse transcription real-time quantitative polymerase chain reaction (RT-qPCR) as the number of viral genomes per ng of total RNA in the plants as described below. Viral load was only estimated after passages 1, 4, 7, 9, 10, and 12 with three-fold replication.

Plant genotypes were clustered according to their phenotypic similarity in response to TuMV infection using the *k*-means method. Goodness of fit of models with increasing number of clusters was evaluated using the minimum value of Bayes information criterion (*BIC*) and the associated Akaike’s weights.

### Experimental evolution

Five TuMV lineages were evolved during 12 consecutive serial passages in each one of the five selected genotypes. To begin the evolution experiment, 10 *A. thaliana* plants per lineage and genotype were inoculated as described above with the virus stock from *N. benthamiana*. The next passages were made by harvesting the symptomatic plants at 12 dpi, preparing the infectious sap as above, and inoculating it into a new batch of 10 plants.

Following the evolution experiment, a cross-infection experiment using all 24 surviving viral lineages was conducted in a single experimental block using 10 plants from each combination of the five plant genotypes utilized in the evolution experiment. Each of the 1,200 plants was monitored throughout 14 dpi, and the severity of the symptoms for each viral lineage-host genotype combination recorded.

### Statistical analyses of disease-related phenotypic traits

Prior to any further analysis, infectivity data were probit-transformed to ensure normality. Transformed infectivity, *AUDPS*, symptoms severity, and *AUSIPS* data were fitted to a MANCOVA model with plant genotype (*G*) as main factor, independent evolution lineages (*L*) nested within *G*, and passage (*t*) as a covariable:

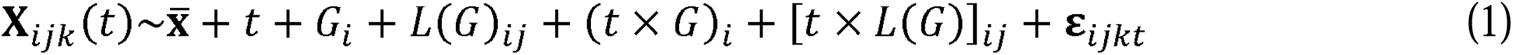

where **X***_ijk_*(*t*) is the vector of disease traits observed at time *t*, for an individual infected plant *k* of evolutionary lineage *j* of genotype *i*, 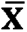 represents the vector of grand mean values and ε*_ijkt_* the vector of errors assumed to be Gaussian. Significance of each factor was tested using the Wilk’s *Λ* summary statistic.

Log-viral load data were fitted to an ANCOVA model with identical structure to equation (1).

Rates of phenotypic evolution for each disease trait were estimated by fitting the corresponding time series to the first-order ARIMA(1,0,0) model:

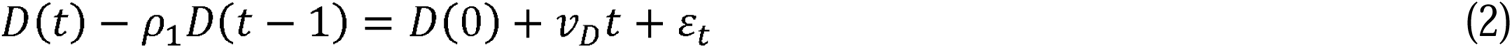

where *D*(*k*) represents the any of the four disease traits at passage *k*, *ρ*_1_ measures the degree of self-similarity in the times-series data, *ε_t_* represents the sampling error at passage *t*, and *v_D_* the linear dependency of trait *D* with *t*, that is, the rate of phenotypic evolution. Estimated rates of phenotypic evolution for the were fitted to a MANOVA with the following structure:

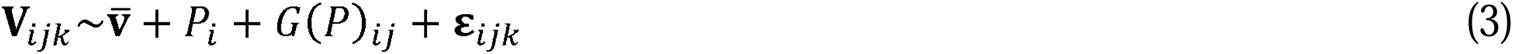

where **V***_ijk_* is the vector of rates of phenotypic evolution, 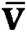 the vector of grand mean values and ε*_ijk_* the vector of Gaussian errors. *P* stands for the epigenetic pathway and *G* is defined as in equation (1).

Cross-infection severity data were fitted to a repeated measures ANOVA in which local host (*LH*) and test host (*TH*) were treated as orthogonal factors, independent evolution lineages (*L*) were nested within *LH* and *t* was treated as the within-plant factor:

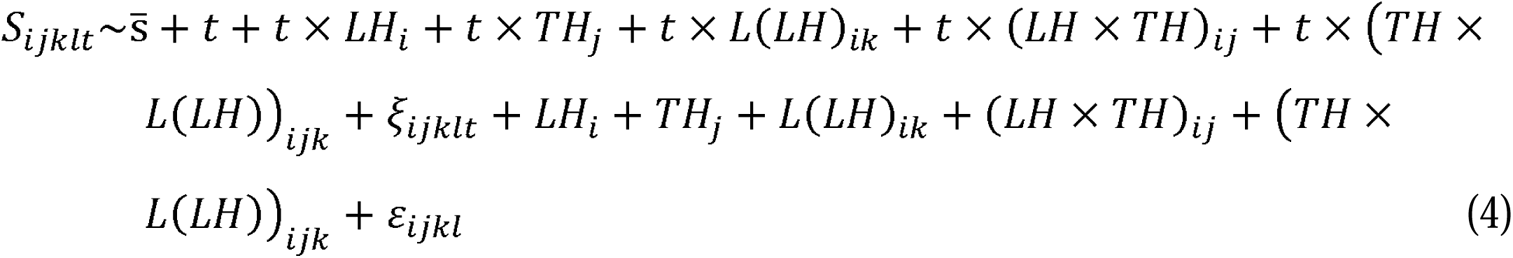

where all terms involving *t* evaluate the within-plant variance, 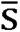 refers to the grand mean value of symptoms severity, *ξ_ijklt_* to the error among observations for a given plant, and *ε_ijkl_* the error among plants, both assumed to be Gaussian.

In all models, the type III sum of squares were used to partition total variance among factors. The magnitude of the effects was evaluated using the 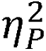 statistic. Values 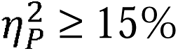 are considered as large effects. All these statistical analyses were done with SPSS version 28.0.1.0 (IBM, Armonk NY, USA).

### Total RNA extractions, quantification of viral load and preparation of samples for RNA-Seq

Pools were made of 10 infected symptomatic plants per lineage, genotype, and serial passage, frozen with liquid N_2_ and preserved at −80 °C until it was homogenized into fine powder. Aliquots of ∼0.1 g were used for total RNA (RNAt) extractions with the Agilent Plant RNA isolation Mini kit (Agilent Technologies, Santa Clara CA, USA). Three aliquots of RNAt per sample were separated and their concentrations adjusted to 50 ng μL^−1^.

Viral load was quantified in each aliquot by RT-qPCR using standard curves and specific forward F117 and reverse F118 primers in an ABI StepOne Plus Real-time PCR System (Applied Biosystems, Foster City CA, USA) as described elsewhere (Cervera et al. 2018; Corrêa et al. 2020; Navarro et al. 2022). For each sample three technical replicates were quantified. Results were analyzed using the StepOne software 2.2.2 (Applied Biosystems).

For RNA-Seq, RNAt from 70 - 90 mg infected and healthy plants was extracted using the GeneJET Plant RNA Purification Mini Kit (Thermo Fisher Scientific, Waltham MA, USA) following the manufacturer’s instructions. RNA quality was checked with NanoDrop One (Thermo Fisher Scientific) and agarose gel electrophoresis and integrity and purity tested with Bioanalyzer 2100 (Agilent Technologies). Libraries preparation and Illumina sequencing was done by Novogene Europe using a NovaSeq 6000 platform and a Lnc-stranded mRNA-Seq library method, ribosomal RNA depletion and directional library preparation, 150 paired end, and 6 Gb raw data per sample. Novogene did quality check of the libraries using a Qubit 4 Fluorometer (Thermo Fisher Scientific), qPCR for quantification and Bioanalyzer for size distribution detection.

### RNA-Seq data processing and measures of viral diversity

The quality of the Fastq files was assessed with FASTQC (Andrews 2010) and MultiQC (Ewels et al. 2016) and preprocessed as paired reads with BBDuk (https://sourceforge.net/projects/bbmap/). Adapters were removed, the first ten 5’ nucleotides of each read were cut, and the 3’ end sequences with average quality below 10 were removed. Processed reads shorter than 80 nucleotides were discarded. The parameter values were set to: *ktrim* = r, *k* = 31, *mink* = 11, *qtrim* = r, *trimq* = 10, *maq* = 5, and *forcetrimleft* = 10, and *minlength* = 80. The processed files were mapped to TuMV isolate YC5 (GenBank, AF530055.2) with the BWA-MEM algorithm (Li 2013). Resulting SAM files were binarized and sorted with SAMtools (Danecek et al. 2021) and duplicates marked with the MarkDuplicates method of GATK version 4.2.2.0 (McKenna et al. 2010).

For TuMV variants calling, the viral assemblies without PCR duplications were used as input for LoFreq (Wilm et al. 2012). Nucleotide counts per position were obtained from the same processed BAM files with pysamstats version 1.1.2 (https://hpc.nih.gov/apps/pysamstats.html). These counts files were used to calculate two diversity metrics: (*i*) Shannon’s entropy per nucleotide site, and (*ii*) Shao et al. (2014) pairwise alignment positional nucleotide counting method which allows calculation of average pairwise distance (*APD*):

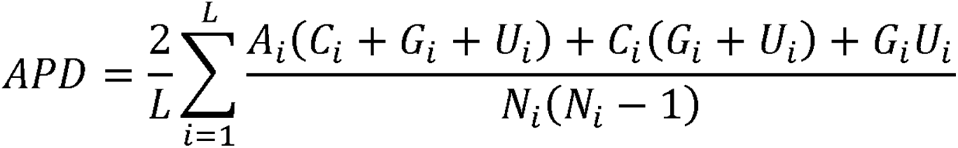

where *L* is the length of the reference genome, *N_i_*the number of total nucleotides and *A_i_*, *C_i_*, *G_i_*, and *U_i_* the counts of each base at site *i*. *APD* data were analyzed using a two-ways non-parametric aligned rank transform (ART) ANOVA (Wobbrock et al. 2011), with passage and plant genotype as orthogonal factors. In this case the type I sum of squares was used. Test was done using the R package ARTool version 0.11.1 (https://cran.r-project.org/web/packages/ARTool/index.html).

Viral consensus sequences were obtained for each lineage after the first and last passages using the consensus function in SAMtools and used to evaluate the possible role of selection in the observed patterns of molecular diversification among lineages evolved in the same and different host genotypes. Four different approaches have been taken. First, we evaluated Nei’s coefficient of nucleotide differentiation *N_ST_* (Nei 1982). Second, Tajima’s *D* test of selection (Tajima 1989) was computed and its significance evaluated using DnaSP version 6.12.03 (Rozas et al. 2017). Third, to evaluate the sign and strength of selection on each of the 10 cistrons in the large ORF, we obtained the average difference among nonsynonymous and synonymous substitution rates, *d_N_* – *d_S_*, using the modified Nei-Gojobori method. These computations were done in MEGA 11 (Tamura et al. 2021); in each analysis the the nucleotide substitution model with the lowest *BIC* was chosen. Standard errors were estimated by 1,000 bootstrap resampling. Finally, we performed a codon selection analysis using the fixed effects likelihood (FEL) method (Kosakovsky Pond and Frost 2005), as implemented in the datamonkey.org server, and with cutoff significance of *P* = 0.05.

The VPg structure was predicted using AlphaFold version 2.3.2 (Collaboratory version: https://colab.research.google.com/github/deepmind/alphafold/blob/main/notebooks/AlphaFold.ipynb#scrollTo=pc5-mbsX9PZC) (Jumper et al. 2021) and visualized with UCSF ChimeraX version 1.2 (Pettersen et al. 2021).

## Results and Discussion

The role of epigenetics in short-term adaptation and long-term evolution has attracted attention as details of the diverse molecular mechanisms involved have been revealed (Rapp and Wendel 2005; Richards et al. 2010; Klironomos et al. 2013; Mendizabal et al. 2014; Stajic et al. 2019; Ashe et al. 2021; Stajic and Jensen 2021). Epigenetic information is more labile than hard-encoded DNA one, which implies higher mutation rates that translate in more phenotypic variance upon which selection may operate facilitating adaptation to sudden challenges (Richards et al. 2010; Banta & Richards 2018). A prototypical example of fast environmental change is created by infection by prevalent parasites (Gómez-Díaz et al. 2012; Kasuga & Gijzen 2013). Of particular interest, viruses have been shown to manipulate the epigenetic pathways during infection. For example, the NS1 protein of influenza A virus contains an amino acid sequence that mimics the host histone H3K4 to hijack transcription elongation factors (Marazzi et al. 2012). Mutant *A. thaliana* plants having loss or gain of epigenetic marks were, respectively, more tolerant and susceptible to TuMV, while infection of wild-type (WT) plants was associated with changes in DNA methylation patterns in protein-coding genes (Corrêa et al. 2020). Together, these observations suggest that epigenetic regulation of host genes impacts viral replication and, therefore, would exert a selective pressure upon evolving viral populations. To test this hypothesis, we have explored the impact that mutations in components of epigenetic regulatory pathways of *A. thaliana* have in TuMV evolution, a virus with high prevalence in this plant (Pagán et al. 2010).

### Classification of plant genotypes based on the TuMV infection phenotype

In a first study we evaluated the effect of mutations in different epigenetic pathways (Table 1) in TuMV infectivity (Fig. S1A), symptom severity (Fig. S1B) and virus accumulation (Fig. S1C).

Fitting the infectivity data in Fig. S1A to a GLM, with plant genotype and days post-inoculation (dpi) as orthogonal factors with a Binomial distribution and probit link function, shows significant differences among plant genotypes that change in magnitude along dpi (test of interaction term: *χ*^2^ = 450.331, 323 df, *P* < 0.001, 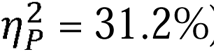). Symptoms severity curves (Fig. S1B) were fitted to a repeated measures ANOVA with dpi as intra-individual factor and plant genotype as inter-individual orthogonal factor. Highly significant differences exist among plant genotypes that depend on the precise time in which individuals are observed (test of interaction term: *F*_247,4927_ = 12.394, *P* < 0.001, 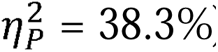). Finally, viral load data (Fig. S1C) were fitted to a GLM with plant genotype and dpi as orthogonal factors with a Gamma distribution and log-link function. Viral load also varies among plant genotypes in a time-dependent manner (test of interaction term: *χ*^2^ = 589.111, 24 df, *P* < 0.001, 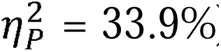). Henceforth, we conclude that significant differences exist among plant epigenetic mutants in their phenotypic response to TuMV infection, ranging from the fast dynamics observed in *rdr1 rdr2 rdr6* to the slow one seen in *ddm1 poliv*.

To cluster plant genotypes on the basis of their phenotypic response to infection, a *k*-means clustering was performed. The smallest *BIC* = 33.9496 (Akaike’s weight 60.22%) was found for *k* = 4 phenogroups (Table S1). The four phenogroups contained a heterogeneous mixture of genotypes. Phenogroup I includes mutants involved in siRNA-dependent RdDM (*ago4*, *ddm1*, *ddm1 poliv*, *ddm1 poliv rdr6*, *ddm1 rdr6*, *polv*, and *rdr6*) and histone modification (*hda6* and *jmj14*) pathways (Table 1). Phenogroup II was formed by a functionally diverse group of mutants: *drm1 drm2*, *poliv*, and *rdr2* affect the siRNA-dependent RdRM, *ibm1* is a H3K9 histone demethylase, and *ros1* is a DNA demethylase (Table 1). Phenogroup III was formed by *dcl2 dcl3 dcl4*, *drm1 drm2 cmt3*, *rdr1 rdr2 rdr6*, and WT. This phenogroup shows functional coherence, since all these mutants affect different steps of the RNA silencing and siRNA-dependent RdDM pathways (Table 1). Finally, phenogroup IV included *atx1* (H3K4 histone methyltransferase) and *cmt3* (unable of maintaining siRNA-independent CHG methylation) (Table 1).

With these findings in hand, we decided to use three mutants involved in siRNA-dependent RdDM (*dcl2 dcl3 dcl4*, *ddm1* and *polv* from phenogroups III, I and I, respectively) and *jmj14* (from phenogroup I) involved in histone demethylation, as hosts for the evolution experiments (Fig. S2). WT plants were included to set the baseline of virus evolution in a host with fully operational epigenetic pathways.

### Evolution of disease-related traits and viral load

The evolution of the four disease-related traits evaluated for each lineage and host genotype are shown in Fig. 1. Though time-series are noisy, in all cases a clear increasing trend has been observed (*i.e*., infections became more virulent), with the exception of lineage *dcl2 dcl3 dcl4*/L3 that got extinct at passage eight. The data were fitted to equation (1) (Table S2). All factors in the model, except lineage, were highly significant (in all cases *P* ≤ 0.005), though the magnitude of the effects varied widely. Net differences among plant genotypes showed the smallest effect (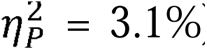), while the largest effect was associated with differences among evolutionary passages (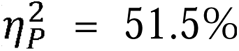). Interestingly, the differences among viral lineages evolved in different plant genotypes varied in a moderate extent along passages (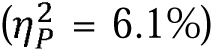). Diversification among viral lineages evolved in the same plant host genotype follow divergent temporal trajectories (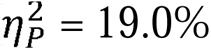).

**Figure 1.**
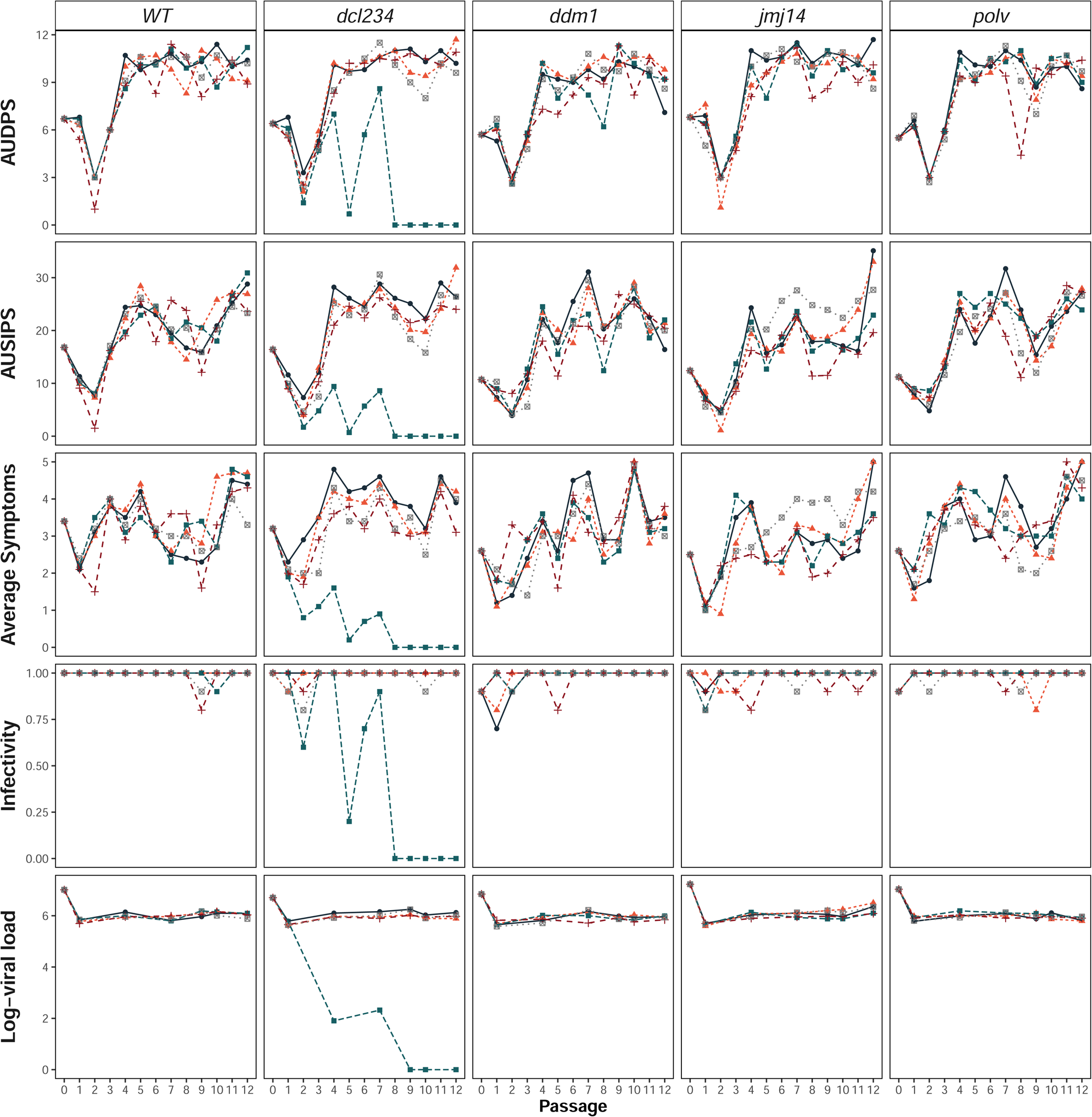
Time-series of the disease-related traits along the evolution experiment for five lineages per plant genotype. Lineages are represented by different colors and symbols. First row: *AUDPS*. Second row: *AUSIPS*. Third row: average symptoms severity measured 14 dpi. Fourth row: infectivity measured at the end of each evolutionary passage (14 dpi). Fifth row: log-viral load (genomes per ng of total RNA).

The last row in Fig. 1 shows the evolution of viral load. Overall, it shows an increase along the evolution experiment, with the exception of the above-mentioned extinct lineage *dcl2 dcl3 dcl4*/L3 and lineage *polv*/L2, which shows a net decrease. Fitting log-viral load data to an ANCOVA similar to equation (1) (Table S3) identifies the same significant factor than for the disease-related traits (all *P* ≤ 0.0004): plant genotype had a very large effect (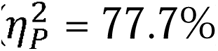), log-viral load increase along evolution passages also in a large magnitude (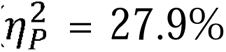) and the magnitude of the observed passage-effect depended on the plant genotype (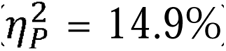), with lineages evolved in a particular plant genotype also differing in their temporal dynamics (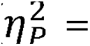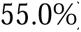). As for the disease-related traits, no net differences exist among lineages evolved in the same plant genotype.

The evolutionary trajectories of the three disease-related traits show a good degree of parallelism, with a sudden reduction after the first passage that is rapidly recovered and further increased after four passages. Thereafter, the rate of evolution slows down to a plateau value. In qualitative terms, this dynamic profile is equivalent for all the host genotype although qualitative differences exist (see below). Such pattern of phenotypic evolution has commonly been observed in evolution experiments with microorganisms (Ebert 1998; Elena and Lenski 2003), and in particular in this experimental pathosystem (Navarro et al. 2022) and can be explained by lineages reaching a (local) maximum in a phenotypic landscape. Interestingly, this observation suggests the existence of a maximum threshold value for the virulence that can be achieved by TuMV in this host. By contrast, either infectivity nor viral load experienced such biphasic shape. Infectivity remained more or less constant and close to the maximum value along the evolution experiment, suggesting no genetic variation in the viral population exists that controls this trait. Viral load experienced a significant reduction after the first passage in all host genotypes and lineages to remain systematically lower than the ancestral strains until the end of the experiment. Nonetheless, an overall net increase from the third passage onwards, has been observed.

Recent studies have linked the evolution of virulence with epigenetic regulation of the pathogen’s genome (Gómez-Díaz et al. 2012; Kasuga & Gijzen 2013). Epigenetic regulation of microbial virulence factors would allow pathogens to quickly accommodate their gene expression to new host species or genotypes. This mechanism, which would be of interest for pathogens with complex genomes such as protozoa, fungi, bacteria or even large DNA viruses that alternate latent with lytic replication cycles [*e.*g., herpes simplex (Singh & Tscharke 2020) or Epstein-Barr (Murata et al. 2021)]. However, direct epigenetic regulation of lytic RNA viruses’ gene expression seems a less efficient mechanism. Instead, RNA viruses manipulate the epigenetic marks of their hosts to improve their own replication (Marazzi et al. 2012; Diezma-Navas et al. 2019; Leone et al. 2020; Corrêa et al. 2020).

### Evolution of the relationship between TuMV accumulation and virulence

Models of virulence evolution typically assume that pathogens’ replication within the host is what causes disease severity (Lenski and May, 1994) and hence a positive association must exist between virulence and accumulation. However, experimental evidence to support this assumption is rather limited for plant-virus pathosystems (Pagán et al. 2007; Agudelo-Romero et al. 2008; Doumayrou et al. 2012). To test this hypothesis, we examined the relationship between log-viral load and the four virulence-related traits. Partial correlation coefficients (controlling for plant genotypes) between log-viral load and the four disease-related traits were computed at different time points during the course of evolution (Fig. S3). In summary, at the beginning of the experiment, viral accumulation was correlated with infectivity (*r_p_* = 0.720, 22 d.f., *P* < 0.001) and *AUDPS* (*r_p_*= 0.698, 22 d.f., *P* < 0.001), but not with average symptoms severity (*r_p_* = 0.145, 22 d.f., *P* = 0.499) nor *AUSIPS* (*r_p_* = 0.207, 22 d.f., *P* = 0.331). In other words, for the ancestral TuMV isolate, viral accumulation was associated with traits related with the appearance of symptoms, but not with their severity. After 12 passages of evolution, however, all four disease-related traits were strongly associated with viral load: infectivity (*r_p_* = 0.990, 22 d.f., *P* < 0.001; representing 37.6% increase in the strength of the association), *AUDPS* (*r_p_* = 0.885, 22 d.f., *P* < 0.001; 26.8% stronger association), symptoms severity (*r_p_* = 0.813, 22 d.f., *P* < 0.001; 460.0% stronger), and *AUSIPS* (*r_p_*= 0.785, 22 d.f., *P* < 0.001; 279.0% stronger).

Most evolution experiments with pathogenic microorganisms show that a selective advantage exists in serial passage experiments for the parasite strain with the highest numerical representation in the transferred inoculum. Within-host host competition and selection for a faster parasite growth rate could be the driving mechanism (Ebert 1998). Our results are relevant in two aspects to this question. Firstly, we found that virus accumulation was associated with increased transmissibility (*i.e*., infectivity and *AUDPS*), and that the association became stronger as the lineages adapted to their local hosts. Secondly, the observation that a significant association between virus accumulation and virulence-related traits (symptoms severity and *AUSIPS*) was built as viral lineages adapted to their local host genotypes suggests that virulence may result for enhanced TuMV replication in the absence of epigenetically-regulated antiviral responses. Interestingly, evolution experiments with other potyvirus, tobacco etch virus (TEV), in different natural accessions of *A. thaliana* with allelic variation in the *RESTRICTED TEV MOVEMENT* (*RTM*) gene, and therefore in their susceptibility to TEV, found significant increases in virus accumulation without changes in virulence (measured as reduction in weight due to infection) (Hillung et al. 2014). This discrepancy between two closely related viruses and different genetic variants of the same host suggests that the association between viral load and virulence shall depend on the specific combination of virus and host genotypes. In particular, virulence would not depend on viral load (*i*) if the extent of damage is not proportional to the amount of viral particles, as in the case of hypersensitive responses (Morel and Dangl 1997), (*ii*) if expressing resistance pathways is costly (Heidel et al. 2004; Traw et al. 2007), (*iii*) if allocating resources to defenses detracts from vegetative growth or reproductive effort (Heil 2001; Pagán et al. 2008), or (*iv*) if the relationship between virus growth rate and virulence is non-monotonic and growth rate itself depends on the metabolic efficiency of the replication process (Lindsay et al. 2023).

### Host epigenetic pathways determine the rates of TuMV phenotypic evolution

To further evaluate the role of mutations in epigenetic pathways in virus evolution, we calculated the rates of phenotypic evolution by fitting the time-series data shown in Fig. 1 to equation (2).

Using the rates of evolutionary change estimated for each lineage as independent replicates, we fitted equation (3). The average rates of phenotypic evolution are displayed in Fig. 2 and the corresponding MANOVA in Table S4. Firstly, significant overall differences between epigenetic pathways exist of very large magnitude 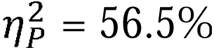). Lineages evolved in WT plants typically, but not always, exhibit the slowest rates of phenotypic evolution. On average, viral lineages evolved in the histone modification mutant *jmj14* exhibit faster rates of phenotypic change than those evolved in RdDM mutants (Fig. 2). Hence, histone modifications seem to exert a stronger selective pressure upon evolving viral populations than changes in DNA methylation profiles. Secondly, mutants within pathways show only marginally significant differences on rates of phenotypic evolution, despite being of large magnitude (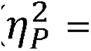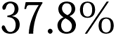). Regardless of the disease phenotype being assessed, viral lineages evolved in *ddm1*, which exhibit compromised siRNA-independent methylation, exhibit the quickest rates of phenotypic change, followed by those evolved in the RdDM mutants *polv* and *dcl2 dcl3 dcl4* (Fig. 2). Notably, the rates of evolution of the five traits are strongly and positively correlated (partial correlation controlling for plant genotype: *r_p_* ≥ 0.750, 22 d.f., *P* < 0.001).

**Figure 2.**
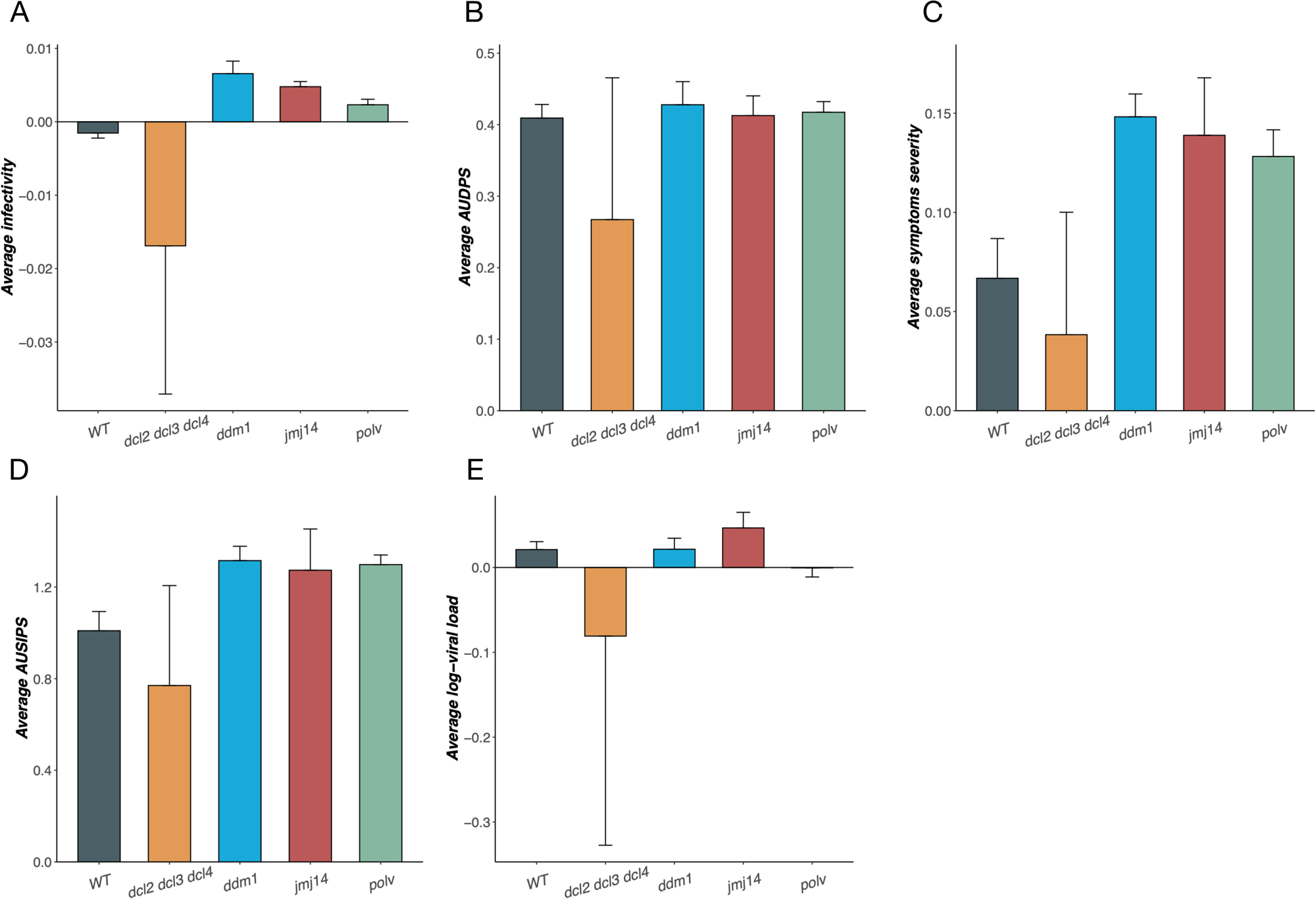
Average rates of evolution for the four disease-related traits measured. Mutants *dcl2 dcl3 dcl4* and *polv* affect RdDM, mutant *ddm1* affect sRNA-independent DNA methylation, while mutant *jmj14* affects histone methylation pathway. (A) Average infectivity, (B) average *AUDPS*, (C) average severity of symptoms, (D) average *AUSIPS*, and (E) log-average viral load. In all cases, error bars represent ±1 SEM.

Previous studies with TEV and TuMV have shown that the rate of evolution inversely correlates with the degree of susceptibility to infection, with faster phenotypic evolution taking place in less susceptible hosts (Hillung et al. 2014; González et al. 2019; Navarro et al. 2022). These previous studies included *A. thaliana* genotypes that differed in mutations affecting specific disease signaling pathways or well-known resistance genes. Here we have extended these observations to the case of mutations in essential components of epigenetically-mediated regulatory pathways. At the one side, our finding that mutations in the H3K4 histone demethylase JMJ14 are associated with the fastest rates of evolution are consistent with the fact that the *jmj14* mutant shows increased H3K4me3 and activation of JMJ14 targets, thus resulting in enhanced expression of resistance genes (Corrêa et al. 2020). At the other side, *dcl2 dcl3 dcl4* plants show compromised DCL-dependent RdDM and RNAi pathways, being unable of mounting an efficient siRNA-mediated antiviral response and becoming highly permissive to viral infection. As a consequence of weaker selection, rates of TuMV evolution measured in these plants are the slowest among all host genotypes tested.

In agreement with the results described here for a plant pathosystem, Mongelli et al. (2022) have done evolution experiments with DCV infecting eight *Drosophila melanogaster* genotypes that carried mutations affecting innate immunity antiviral defense pathways. Firstly, they also found that the extent and evolutionary dynamics of viral accumulation and infectivity depended on the particular host genotype, with a clear tendency to increase viral load in immunity-deficient flies. Secondly, also in agreement with our observations, DCV results also highlight pleiotropy (viral genotype-by-host genotype interaction) as a major determinant of viral evolution. Indeed, they clearly show that the fitness effect of mutations that already existed in the standing genetic variation of the stock DCV population was strongly dependent on both, the fly’s genetic background (the same mutation being beneficial or deleterious in different backgrounds) and the virus’ genetic background (epistatic interactions with other mutations in the same haplotype).

### Analysis of cross-infection data

The gene-for-gene and mutation accumulation models represent the two ends of a continuum of possible outcomes of host - pathogen interaction (Agrawal and Lively 2002). With a pure gene-for-gene interaction, the susceptible host types are expected to disappear and the resistant types are expected to dominate the population. *Vice versa*, the most virulent virus allele would become fixed at the cost of milder ones. However, constitutive activation of defenses is known to be costly for *A. thaliana* [*e.g.*, SA-related defense responses pay a fitness cost in absence of pathogens (Traw et al. 2007)] and high virulence usually comes with a cost in terms of pathogen’s transmission (Acevedo et al. 2019). Hence, a pure gene-for-gene strategy seems unlikely to be achieved. By contrast, with a pure mutation accumulation interaction, negative frequency-dependent selection emerges, such that rare *A. thaliana* resistance alleles have advantage and, as a result, a genetic polymorphism shall be maintained (Schmid-Hempel 2011). Symptom progression was evaluated for each one of the 24 evolved TuMV lineages across all five host genotypes. To test the extent of the specificity of TuMV adaptation to their local host genotypes, we fitted the cross-infection data to equation (4). Focusing in the analyses of the inter-plant factors (Table 2), a first highly significant effect of large magnitude (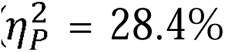) has been observed among different local host genotypes [*LH* in equation (4)], supporting the idea that mutations affecting different epigenetic regulatory pathways determine the evolutionary fate of viruses. A second highly significant effect, yet of moderate magnitude (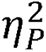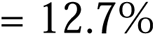), has been observed for the genotype in which viral lineages have been tested [*TH* in equation (4)]. This describes the effect of differences among plant genotypes irrespective of the infecting viral lineage and its past evolutionary history. The interaction between these two main factors [*TH*×*LH* in equation (4)] is also highly significant, suggesting that disease progression associated with the infection of a virus evolved in a given local host depends on the test host genotype; however, the magnitude of this effect is small (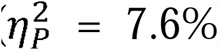). A fourth highly significant effect, though of small magnitude (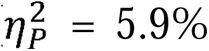), has been observed among viral lineages evolved in the same local host genotype [*L*(*LH*) in equation (4)], which suggests that differences among lineages that have been evolved in the same host genotype are minor compared to differences among lineages evolved in different local hosts. In this sense, these significant effects confirm that infection with some viral lineages generally results in more severe disease in a local host-dependent manner. Finally, the highly significant and of large magnitude (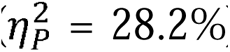) effect associated with the viral lineage by host genotype interaction [*TH*×*L*(*LH*) in equation (4)] confirms that the outcome of infection ultimately depends upon the particular combination of host and viral genotypes. The analyses of the intra-plant factors, drive to identical conclusions and confirm that the severity of symptoms increases along time [*t* in equation (4); 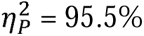].

**Table 2.** Results of fitting the severity of symptoms progression data to the repeated measures ANOVA described in equation (4).

| Effect | SS<sup>1</sup> | d.f. | F | P | $\eta_p^2$ | 1 - $\beta$ |
| --- | --- | --- | --- | --- | --- | --- |
| **Intra-plants tests<sup>2</sup>** |  |  |  |  |  |  |
| dpi (*t*) | 16642.8358 | 5.2755 | 8169.3272 | < 0.0001 | 0.8832 | 1 |
| dpi by local host (*t* × *LH*) | 296.3525 | 21.1018 | 36.3670 | < 0.0001 | 0.1187 | 1 |
| dpi by test host (*t* × *TH*) | 125.5401 | 21.1018 | 15.4057 | < 0.0001 | 0.0540 | 1 |
| dpi by local host by test host (*t* × *LH* × *TH*) | 161.8922 | 84.4073 | 4.9667 | < 0.0001 | 0.0685 | 1 |
| dpi by lineage within local host (*t* × *L(LH)*) | 97.4343 | 105.5091 | 2.3913 | < 0.0001 | 0.0424 | 1 |
| dpi by test host by lineage within local host (*t* × *TH* × *L(LH)*) | 536.7017 | 395.6590 | 3.5126 | < 0.0001 | 0.1961 | 1 |
| Error within dpi | 2200.2133 | 5697.4902 |  |  |  |  |
| **Inter-plants tests** |  |  |  |  |  |  |
| Intersection ($\Sigma$) | 20676.4419 | 1 | 22744.8162 | < 0.0001 | 0.9547 | 1 |
| Local host (*LH*) | 388.7266 | 4 | 106.9032 | < 0.0001 | 0.2836 | 1 |
| Test host (*TH*) | 142.2224 | 4 | 39.1124 | < 0.0001 | 0.1265 | 1 |
| Local host by test host (*LH* × *TH*) | 80.3478 | 16 | 5.5241 | < 0.0001 | 0.0756 | 1 |
| Lineage within local host (*L(LH)*) | 61.4782 | 20 | 3.3814 | < 0.0001 | 0.0589 | 1 |
| Test host by lineage within local host (*TH* × *L(LH)*) | 385.9958 | 75 | 5.6615 | < 0.0001 | 0.2822 | 1 |
| Error | 981.7867 | 1080 |  |  |  |  |

<sup>1</sup>Type III sum of squares.
<sup>2</sup>Greenhouse-Geisser correction for data sphericity ( $\epsilon = 0.3768$ ).

These results align with those by Hillung et al. (2014), González et al. (2019) and Navarro et al. (2022), all pointing towards the evolution of a combination of specialist and generalist viral lineages and of more permissive and resistant host genotypes [significant *TH*×*LH* and *TH*×*L*(*LH*) in equation (4)]. Indeed, these studies have also shown that more permissive hosts *dcl2 dcl3 dcl4* selected for more specialized viruses while more restrictive hosts *jmj14* selected for more generalist viruses, hence matching the expectation from the gene-for-gene model.

### Genomic changes in evolved TuMV lineages

Fig. 3A shows the change in viral genetic diversity within lineages, measured as *APD*, experienced by each lineage in the course of evolution. In all cases, a significant increase has been observed (*F*_1,19_ = 165.788, *P* < 0.001), being on average 30.2% ±2.9 larger after 12 passages. More interestingly, significant differences in the amount of genetic diversity accumulated in the lineages exist among plant genotypes (*F*_4,19_ = 3.369, *P* = 0.030). Indeed, the magnitude of the differences among lineages evolved in different plant genotypes depended on the passage being characterized (*F*_4,19_ = 3.884, *P* = 0.018). The average magnitude of *APD* increases ranged from the 20.0% ±4.8 observed for lineages evolved in *ddm1* to the 44.7% ±5.0 observed among lineages evolved in *dcl2 dcl3 dcl4*, with lineages evolved in WT plants showing the second smaller increase in genetic variability.

**Figure 3.**
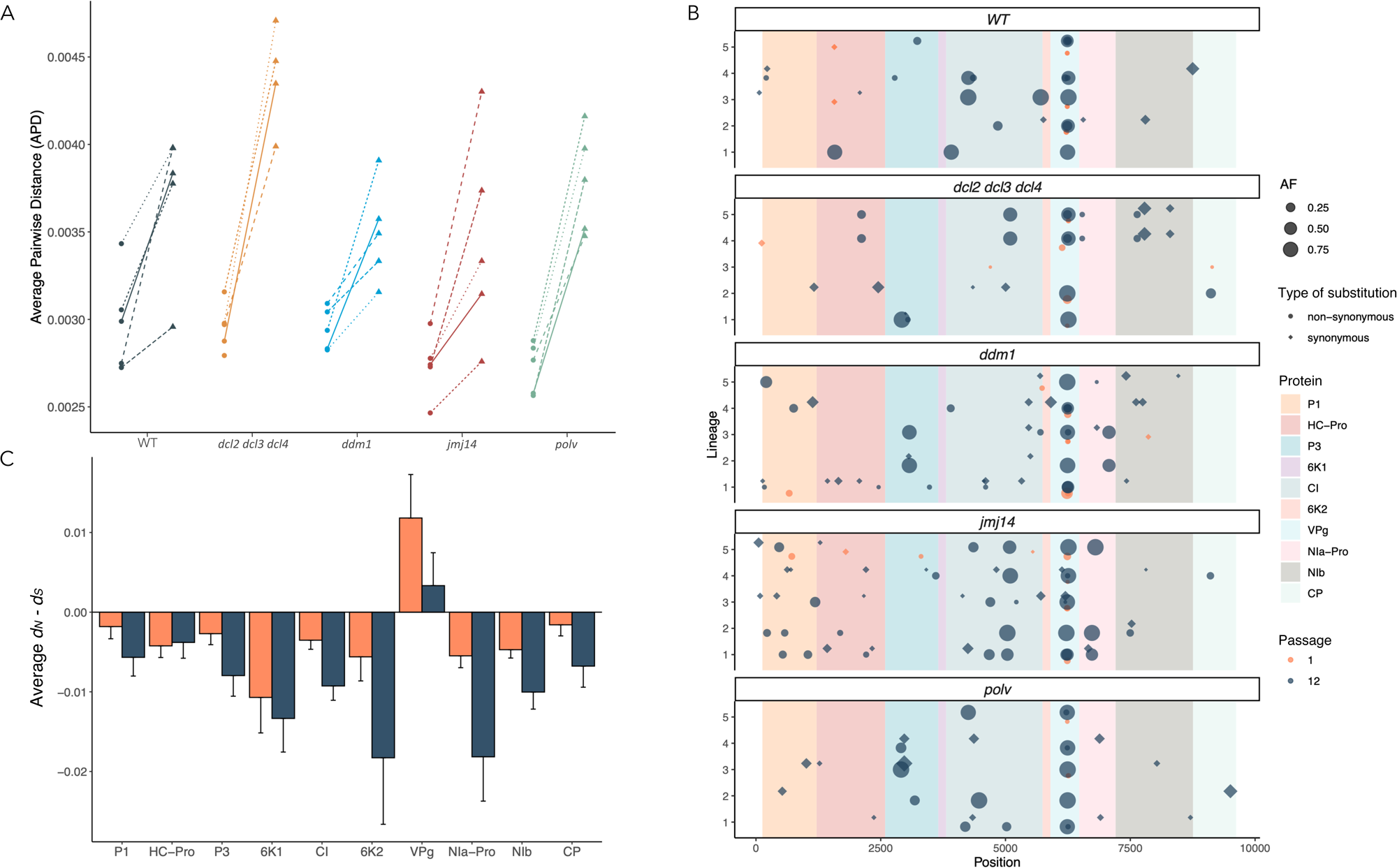
(A) Observed mean change in viral diversity between the first and last evolution passages. Diversity is measured as average pairwise *p*-distances (*APD*). Different lineages are represented with different line types. *APD* at passage 1 is represented by solid circles while solid triangles represent *APD* after passage 12. Different colors correspond to different plant genotypes. (B) Distribution of variability along TuMV genome observed for each lineage and host genotype. In blue, SNVs observed after passage 12, in orange variants observed after passage 1. The size of the balls is proportional to allele frequency (> 10% in all cases). (C) Average *d_N_* – *d_S_*per cistron. Orange bars represent passage 1 and dark blue bars passage 12. Error bars correspond to ±1 SD.

Fig. 3B summarizes the distribution of genetic diversity along the genomes of the evolved lineages. For the shake of a better visualization, only single nucleotide variants with an allele frequency > 10% are shown, although all variants identified (Fig. S3) would be used in the analyses below. The total number of segregating sites after passage 1 was 518, rising up to 1,206 (2.33-fold increase) by passage 12. Interestingly, most of the segregating sites at both times were singletons (425 after passage 1 and 1,038 after passage 12). Treating each independent lineage as an observation and each host genotype as a subpopulation, the average nucleotide diversity within-host genotypes observed was *π_S_*(1) = (3.869 ±0.426) ×10^−3^, increasing to *π_S_*(12) = (8.173 ±0.060) ×10^−3^ (2.11 times more variability). The nucleotide diversity for the entire sample was *π_T_*(1) = (3.879 ±0.409) ×10^−3^ and increased to *π_T_*(12) = (8.639 ±0.637) ×10^−3^ (2.23 times more). Hence, the estimated interhost genotype nucleotide diversities at both time points were *δ_ST_*(1) = (0.965 ±4.137) ×10^−5^ and *δ_ST_*(12) = (4.658 ±0.870) ×10^−4^, which represents a 48.3-fold increase in differences among viral lineages evolved in different host genotypes. These figures can be combined to compute the coefficient of nucleotide differentiation (Nei 1982) among host genotypes at the beginning of the experiment [*N_ST_*(1) = (0.249 ±1.054) ×10^−2^, *z* = 0.236, *P* = 0.407)] and at the end [*N_ST_*(12) = (5.392 ±0.087) ×10^−2^, *z* = 61.977, *P* < 0.001]. Hence, we conclude that genetically homogeneous founding viral populations had generated significant differentiation after 12 passages of evolution in different host genotypes (significant difference between both *N_ST_*values: *z* = 4.862, *P* < 0.001).

### Assessing the role of selection upon TuMV genome

To assess whether selection played a role in virus genetic differentiation among plant genotypes, we performed a Tajima’s *D* test (Tajima 1989) on the consensus sequences per sample and found it was significantly negative (*D* = −2.457, *P* = 0.006) after passage 1, becoming further more by the end of the experiment (*D* = −2.567, *P* < 0.001). These negative *D* values are compatible with the action of purifying selection, the presence of segregating slightly deleterious mutations or fast population expansions (Yang 2006). It is expected that in fast expanding populations, many new mutations may rise in frequency, thus being observed as singletons in each lineage. Singletons inflate the number of segregating sites and thus cause *D*D< 0. Indeed, this was the case here: 82.0% of the observed variable sites were singletons after passage 1, and 86.1% after passage 12, thus the observed pattern of molecular diversity among lineages evolved in the same and in different host genotypes was compatible with the fixation of mutations during the exponential growth experienced by viral populations during the systemic colonization of plants after each passage.

The three above mentioned explanations for a negative *D* value are not mutually exclusive; hence, room for natural selection still exists. Therefore, we sought to evaluate the role of selection at the cistron level. To do so, we computed the average *d_N_* – *d_S_* for each cistron in the main ORF at both passages (Fig. 3C). Overall, nine cistrons were, on average, under purifying selection, which became more intense by the end of the evolution experiment. This is particularly relevant for NIb (Fig. S4), which shows two highly variable regions in passage 1, one of which is systematically homogenized, in all host genotypes except in *polv*, by the end of the experiment. The only cistron that showed signatures of positive diversifying selection was *VPg*, though the strength of this selection also varied along the evolution experiment, being stronger at the beginning and weaker by the end. To further expand this observation, we performed a FEL codon-based selection analysis (Fig. 4A). Codons encoding for G3, Y25 and P78 have been found to be under significant purifying selection. The mutation at Y25 was observed at the two sequenced time points, while the other two mutations were seen only in passage 12. Nine nonsynonymous mutations have been diagnosed as under significant positive diversifying selection. In codon 79, mutations L79F and L79I were observed in different lineages, regardless of the host genotype, after the first passage. However, by the end of the experiment only the ancestral allele L was retained in all 24 lineages. The other eight nonsynonymous mutations all appeared in a specific domain of VPg that encompasses amino acids 110 – 120 (Fig. 4B). All these polymorphisms show a remarkably similar pattern: most lineages show alternative alleles after the first passage of evolution. However, after the last passage, many of these mutations had reverted to the corresponding ancestral alleles. Let’s take codon 115 as an illustrative example (Fig. 4A), in all cases, after passage 1 all lineages showed different new alleles [*e.g*., N115A (in 4 lineages), N115D (1), N115G (12), N115L (1), N115P (2), N115Q (1), and N115R (4)]. However, by the end of the experiment, the ancestral allele N was fixed back in a majority of lineages (13), while the rest shared some new alleles [*e.g*., N115D (*ddm1*/L1 and WT/L1), N115G (*ddm1* lineages L2 and L3) and N115S (*jmj14* lineages L1 and L4, *polv* lineages L2, L3 and L4 and WT lineages L3 and L4)]. Together, this observation of transient rise in frequency of novel alleles followed by a reversal to ancestral ones is in agreement with the expectation of pervasive leap-frog events (Gerrish and Lenski 1998). According to this effect, the majority of genotypes at some time point is less closely related to the immediately preceding majority genotype than to an earlier one. Given that reversion mutations are unlikely, the permanence of ancestral alleles at very low frequency right after the switch from WT to mutant host genotypes seems a more parsimonious explanation for the leap-frogs.

**Figure 4.**
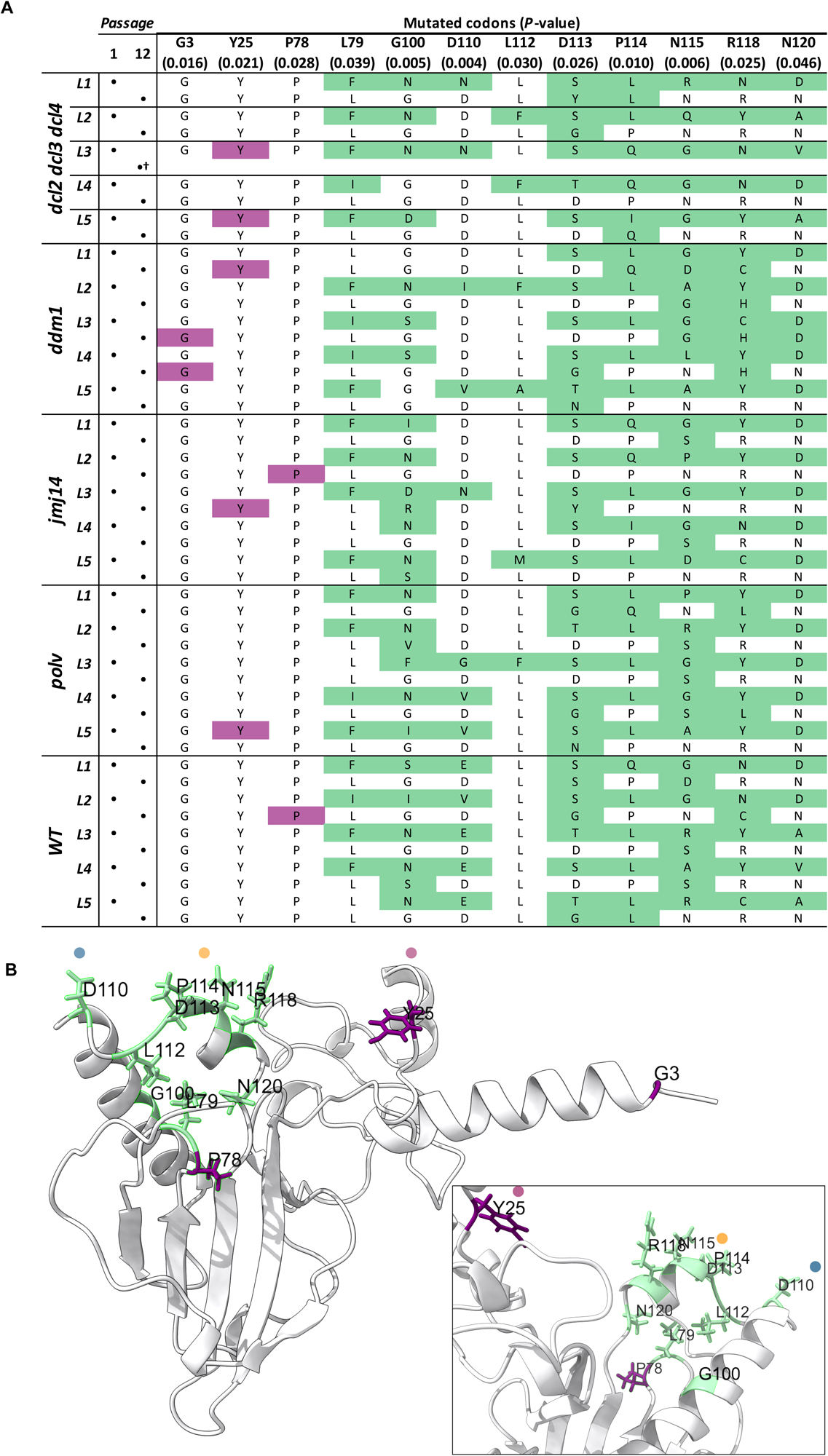
(A) Results of the FEL analysis showing positions at the *VPg* cistron under significant negative stabilizing selection (purple) and positive diversifying selection (green). Rows correspond to the different lineages are paired by sequenced passages. Each cell contains the observed amino acid. The header shows the reference amino acid and the probability of accepting the neutral evolution hypothesis provided by FEL. (B) VPg structure predicted with AlphaFold The position of the amino acids under selection are shown with the same colors as in panel A on the protein structure. The inset shows the domain encompassing amino acids 110 - 120. The blue, yellow and purple dots have been added to facilitate orientation between the two views.

No obvious association between the alleles fixed at the end of the evolution experiment and host genotypes exists, except perhaps for allele R118H (keeping a positively charged side chain and neutral functional effect) which has been found only in three lineages evolved in *ddm1*. Interestingly, this mutation was also observed by Navarro et al. (2022) associated with TuMV lineages evolved in jasmonate insensitive mutant plants showing an enhanced systemic acquired resistance.

*VPg* has been commonly found as a target of selection in evolution experiments of potyviruses in *A. thaliana* (Agudelo-Romero et al. 2008; Gallois et al. 2010; Hillung et al. 2014; González et al. 2019; González et al. 2021; Navarro et al. 2022; Melero et al. 2023). VPg plays many essential roles in genome transcription (it is linked to the 5’-end of the viral genome and provides the hydroxyl group that primes the synthesis of the complementary strains by the viral RdRp; Murphy et al. 1996), translation [directly interacts with the eukaryotic initiation factors eIF(iso)4E and eIF(iso)4G; Léonard et al. 2000], and interacts with all other viral proteins and some of the host cell proteins (Martínez et al. 2023). In addition, the role of VPg as a target of selection in crops carrying resistance genes has been amply demonstrated. One of the most extensively used resistance genes against potato virus Y in commercial pepper cultivars is *pvr2*, which has many different alleles (Nicaise et al. 2007; Charron et al. 2008). The *pvr2* locus encodes for the eIF4E factor that. Interestingly, all the resistance-breaking viral isolates found so far contain mutations in the *VPg* cistron (Duprat et al. 2002; Moury et al. 2004; Ayme et al. 2006).

## Conclusions

We have shown that epigenetic regulatory pathways not only have an effect in the antiviral responses of hosts and that viruses interfere with such defenses, but also that epigenetic regulation impacts the evolution of RNA viruses. Previous studies have shown that mutations in resistance/susceptibility recessive genes and in dominant genes involved in defense responses to infection modulate the evolution of viral populations. A common observation of these experiments is that more permissive hosts select for less virulent pathogens, while more virulent pathogens are selected in more resistant hosts. Here we extend these previous results to the case of epigenetic regulation of defense genes. In particular, we describe how mutations in the two main epigenetic pathways (*i.e*., DNA methylation and histone modification) condition the evolution of a plant RNA virus. Parallelizing previous results, more resistant genotypes, in this particular case a mutant affecting histone modification, selected for more virulent viruses, while a mutant unable of producing siRNAs and thus particularly sensitive to infection, selected for less virulent viral strains. Likewise, we confirm that VPg is a main target of selection also in the context of a potyviral interplay with the host epigenetic regulation, which suggests a possible role of this multifunctional viral protein in manipulating unknown aspects of the host response to infection. In a follow-up manuscript, we will tackle this issue by deeply characterizing host genotype-specific transcriptomic responses to infection with the ancestral and evolved TuMV lineages.

## Author contributions

S.F.E. conceived the study. S.A. performed the experiments. M.J.O.U., R.L.C. and S.F.E. analyzed the data. S.F.E. drafted the initial version of the manuscript. All authors contributed to later versions of the manuscript.

## Acknowledgements

We thank Francisca de la Iglesia and Paula Agudo for excellent technical support and the EvolSysVir lab members for comments and fruitful discussions. This work was supported by grant PID2019-103998GB-I00 funded by MCIN/AEI/10.13039/501100011033 and by Generalitat Valenciana grant PROMETEO/2019/012 to S.F.E.

## Supplementary material

Supplementary material is available online at *Evolution Letters* (https://academic.oup.com/evlett/****).

## Data availability

Raw Illumina RNA-seq data generated for this study are available in NCBI SRA under BioProject accession XXX. All R scripts developed in this work are available at https://github.com/MJmaolu/evolution_TuMV_in_A.thaliana_Epigenetic_Mutants/tree/main.

## References

Acevedo, M.A., Dillemuth, F.P., Flick, A.J., Faldyn, M.J., & Elderd, B.D. (2019) Virulence-driven trade-offs in disease transmission: a meta-analysis. Evolution, 73, 636–647.

Agrawal, A., & Lively, C. (2002) Infection genetics: gene-for-gene versus matching-alleles models and all points in between. Evol. Ecol. Res., 4, 79–90.

Agudelo-Romero, P., de la Iglesia, F., & Elena, S.F. (2008) The pleiotropic cost of host-specilization in tobacco etch potyvirus. Infect. Genet. Evol. 8, 806–814.

Álvarez-Venegas, R., Pien, S., Sadder, M., Witmer, X., Grossniklaus, U. & Avramova, Z. (2003) ATX-1, an Arabidopsis homolog of trithorax, activates flower homeotic genes. Curr. Biol., 13, 627–637.

Andrews, S. (2010) FASTQC: a quality control tool for high throughput sequence data. Available online at http://www.bioinformatics.babraham.ac.uk/projects/fastqc/.

Ashe, A., Colot, V., & Olroyd, B.P. (2021) How does epigenetics influence the course of evolution? Phil. Trans. R. Soc. B, 376, 20200111.

Ayme, V., Souche, S., Caranta, C., Jacquemond, M., Chadoeuf, J., Palloix, A., et al. (2006) Different mutations in the genome-linked protein VPg of potato virus Y confer virulence on the *pvr2^3^* resistance in pepper. Mol. Plant Microbe Interact., 19, 557–563.

Banta, J.A., & Richards, C.L. (2018) Quantitative epigenetics and evolution. Heredity, 121, 210–224.

Böhmdorfer, G., Sethuraman, S., Rowley, M.J., Krzyszton, M., Rothi, M.H., Bouzit, L., & Wierzbicki, A.T. (2016) Long non-coding RNA produced by RNA polymerase V determines boundaries of heterochromatin. eLife, 5, e19092.

Bond, D.M., & Baulcombe, D.C. (2014) Small RNAs and heritable epigenetic variation in plants. Trends Cell Biol., 24, 100–107.

Borges, F. & Martienssen, R.A. (2015) The expanding world of small RNAs in plants. Nat. Rev. Mol. Cell. Biol., 16, 727–741.

Bouché, N., Lauressergues, D., Gasciolli, V. & Vaucheret, H. (2006) An antagonistic function for Arabidopsis DCL2 in development and a new function for DCL4 in generating viral siRNAs. EMBO J., 25, 3347–3356.

Boyes, D.C., Zayed, A.M., Ascenzi, R., McCaskill, M.J., Hoffman, N.E., Davis, K.R., et al. (2001) Growth stage-based phenotypic analysis of Arabidopsis: a model for high throughput functional genomics in plants. Plant Cell, 13, 1499–1510.

Butković, A., González, R., Rivarez, M.P.S. & Elena, S.F. (2021) A genome-wide association study identifies *Arabidopsis thaliana* genes that contribute to differences in the outcome of infection with two turnip mosaic potyvirus strains that differ in their evolutionary history and degree of host specialization. Virus Evol., 7, veab063.

Cao, X. & Jacobsen, S.E. (2002) Role of the Arabidopsis *DRM* methyltransferases in *de novo* DNA methylation and gene silencing. Curr. Biol., 12, 1138–1144.

Carr, J.P., Lewsey, M.G. & Palukaitis, P. (2010) Signaling in induced resistance. Adv. Virus Res., 76, 57–121.

Cervera, H., Ambrós, S., Bernet, G.P., Rodrigo, G. & Elena, S.F. (2018) Viral fitness correlates with the magnitude and direction of the perturbation induced in the host’s transcriptome: the tobacco etch potyvirus - tobacco case study. Mol. Biol. Evol., 35, 1599–1615.

Chan, S.W.L., Henderson, I.R., Zhang, X., Shah, G., Chien, J.S.C. & Jacobsen, S.E. (2006) RNAi, DRD1, and histone methylation actively target developmentally important non-CG DNA methylation in Arabidopsis. PLoS Genet., 2, e83.

Charron, C., Nicolaï, M., Gallois, J.L., Robaglia, C., Moury, B., Palloix, A., et al. (2008) Natural variation and functional analyses provide evidence for co-evolution between plant eIF4E and potyviral VPg. Plant J., 54, 56–68.

Chen, C.C., Chao, C.H., Chen, C.C., Yeh, S.D., Tsai, H.T. & Chang, C.A. (2003) Identification of turnip mosaic virus isolates causing yellow stripe and spot on calla lily. Plant Dis., 87, 901–905.

Corrêa, R.L., Sanz-Carbonell, A., Kogej, Z., Müller, S.Y., Ambrós, S., López-Gomollón, S., et al. (2020) Viral fitness determines the magnitude of transcriptomic and epigenomic reprograming of defense responses in plans. Mol. Biol. Evol., 37, 1866–1881.

Cuerda-Gil, D. & Slotkin, R.K. (2016) Non-canonical RNA-directed DNA methylation. Nat. Plants, 2, 16163.

Danecek, P., Bonfield, J.K., Liddle, J., Marshall, J., Ohan, V., Pollard, M., et al. (2021) Twelve years of SAMtools and BCFtools. GigaScience, 10, giab008.

Diezma-Navas, L., Pérez-González, A., Artaza, H., Alonso, L., Caro, E., Llave, C., et al. (2019) Crosstalk between epigenetic silencing and infection by tobacco rattle virus in Arabidopsis. Mol. Plant Pathol., 20, 1439–1452.

Doumayrou, J., Avellan, A., Froissart, R., & Michalakis, Y. (2012) An experimental test of the transmission-virulence trade-off hypothesis in a plant virus. Evolution, 67, 477–486.

Dowen, R.H, Pelizzola, M., Schmitz, R.J., Lister, R., Dowen, J.M., Nery, J.R., et al. (2012) Widespread dynamic DNA methylation in response to biotic stress. Proc. Natl. Acad. Sci. USA, 109, E2183–E2191.

Duprat, A., Caranta, C., Revers, F., Menand, B., Browning, K.S., & Robaglia, C. (2002) The Arabidopsis eukaryotic initiation factor (iso)4E is dispensable for plant growth but required for susceptibility to potyviruses. Plant J., 32, 927–934.

Earley, K., Lawrence, R.J., Pontes, O., Reuther, R., Enciso, A.J., Silva, M., et al. (2006) Erasure of histone acetylation by Arabidopsis *HDA6* mediates large-scale gene silencing in nucleolar dominance. Genes Develop., 20, 1283–1293.

Ebert, D. (1998) Experimental evolution of parasites. Science, 282, 1432–1435.

Elena, S.F., & Lenski, R.E. (2003) Evolution experiments with microorganisms: the dynamics and genetic bases of adaptation. Nat. Rev. Genet., 4, 457–469.

Ewels, P., Magnusson, M., Lundin, S. & Käller, M. (2016) MultiQC: Summarize analysis results for multiple tools and samples in a single report. Bioinformatics, 32, 3047–3048

Gallois, J.L., Charron, C., Sánchez, F, Pagny, G., Houvenaghel, M.C., Moretti, A., et al. (2010) Single amino acid changes in the turnip mosaic virus viral genome-linked protein (VPg) confer virulence towards *Arabidopsis thaliana* mutants knocked out for eukaryotic initiation factors eIF(iso)4E and eIF(iso)4G. J. Gen. Virol. 91, 288–293.

Garcia-Ruiz, H., Takeda, A., Chapman, E.J., Sullivan, C.M., Fahlgren, N., Brempelis, K.J., et al. (2010) Arabidopsis RNA-dependent RNA polymerases and Dicer-like proteins in antiviral defense and small interfering RNA biogenesis during turnip mosaic virus infection. Plant Cell, 22, 481–496.

Gerrish, P.J., & Lenski, R.E. (1998) The fate of competing beneficial mutations in an asexual population. Genetica, 102–103, 127-144.

Gómez-Díaz, E., Jordà, M., Peinado, M.A., & Rivero, A. (2012) Epigenetics of host-pathogen interactions: the road ahead and the road behind. PLoS Pathog., 8, e1003007.

Gong, Z., Morales-Ruiz, T., Ariza, R.R., Roldán-Arjona, T., David, L., & Zhu, J.K. (2002) *ROS1*, a repressor of transcriptional gene silencing in Arabidopsis, encodes a DNA glycosylase/lyase. Cell, 111, 803–814.

González, R., Butković, A., & Elena, S.F. (2019) Role of host genetic diversity for susceptibility-to-infection in the evolution of virulence of a plant virus. Virus Evol., 5, vez024.

González, R., Butković, A., Escaray, F.J., Martínez-Latorre, J., Melero, I., Pérez-Parets, E., et al. (2021) Plant virus evolution under strong drought conditions results in a transition from parasitism to mutualism. Proc. Natl. Acad. Sci. USA, 118, e2020990118.

Heil, M. (2001) The ecological concept of costs of induced systemic resistance (ISR). Eur. J. Plant Pathol., 107, 137–146.

Heidel, A.J., Clarke, J.D., Antonovics, J., & Dong, X. (2004) Fitness costs of mutations affecting the systemic acquired resistance pathway in *Arabidopsis thaliana*. Genetics, 168, 2197–2206.

Herr, A.J., Jensen, M.B., Dalmay, T., & Baulcombe, D.C. (2005) RNA polymerase IV directs silencing of endogenous DNA. Science, 308, 118–120.

Hillung, J., Cuevas, J.M., Valverde, S., & Elena, S.F. (2014) Experimental evolution of an emerging plant virus in host genotypes that differ in their susceptibility to infection. Evolution, 68, 2467–2480.

Hung, Y.H., & Slotkin, R.K. (2021) The initiation of RNA interference (RNAi) in plants. Curr. Opin. Plant Biol., 61, 102014.

Jeddeloh, J.A., Stokes, T.L., & Richards, E.J. (1999) Maintenance of genomic methylation requires a SWI2/SNF2-like protein. Nat. Genet., 22, 94–97.

Jumper, J., Evans, R., Pritzel, A., Green, T., Figurnov, M., Ronneberger, O., et al. (2021) Highly accurate protein structure prediction with AlphaFold. Nature, 596, 583–589.

Kachroo, P., Chandra-Shekara, A.C., & Klessig, D.F. (2006) Plant signal transduction and defense against viral pathogens. Adv. Virus Res., 66, 161–191.

Kasuga, T., & Gijzen, M. (2013) Epigenetics and the evolution of virulence. Trends Microbiol., 21, 575–582.

Klironomos, F.D., Berg, J., & Collins, S. (2013) How epigenetic mutations can affect genetic evolution: model and mechanisms. Bioessays, 35, 571–578.

Kosakovsky Pond, S.L., & Frost, S.D.W. (2005) Not so different after all: a comparison of methods for detecting amino acid sites under selection. Mol. Biol. Evol., 22, 1208–1222.

Le, T.N., Schumann, U., Smith, N.A., Tiwari, S., Au, P.C.K., Zhu, Q.H., et al. 2014. DNA demethylases target promoter transposable elements to positively regulate stress responsive genes in Arabidopsis. Genome Biol., 15, 458.

Lenski, R.E., & May, R.M. (1994) The evolution of virulence in parasites and pathogens: reconciliation between two competing hypotheses. J. Theor. Biol., 169, 253–265.

Léonard, S., Plante, D., Wittmann, S., Daigneault, N., Fortin, M.G., & Laliberté, J.F. (2000) Complex formation between potyvirus VPg and translation eukaryotic initiation factor 4E correlates with virus infectivity. J. Virol., 74, 7730–7737.

Leone, M., Zavallo, D., Venturuzzi, A., & Asurmendi, S. (2020) RdDM pathway components differentially modulate *Tobamovirus* symptom development. Plant Mol. Biol., 104, 467–481.

Li, H. (2013) Aligning sequence reads, clone sequences and assembly contigs with BWA-MEM. arXiv. http://arxiv.org/abs/1303.3997.

Liu, P., Cuerda-Gil, D., Shahid, S., & Slotkin, R.K. (2022) The epigenetic control of the transposable element life cycle in plant genomes and beyond. Annu. Rev. Genet., 56, 63–87.

Lindroth, A.M., Cao, X., Jackson, J.P., Zilberman, D., McCallum, C.M., Henikoff, S., et al. 2001. Requirement of *CHROMOMETHYLASE3* for maintenance of CpXpG methylation. Science, 292, 2077–2080.

Lloyd, J.P.B., & Lister, R. (2021) Epigenome plasticity in plants. Nat. Rev. Genet., 23, 55–68.

Loebenstein, G. (2009) Local lesions and induced resistance. Adv. Virus Res., 75, 73–117.

López, A., Ramírez, V., García-Andrade, J., Flors, V., & Vera, P. (2011) The RNA silencing enzyme RNA polymerase V is required for plant immunity. PLoS Genet., 7, e1002434.

López-Gomollon, S., & Baulcombe, D.C. (2022) Roles of RNA silencing in viral and non-viral plant immunity and in the crosstalk between disease resistance systems. Nat. Rev. Mol. Cell Biol., 23, 645–662.

López Sánchez, A., Stassen, J.H.M., Furci, L., Smith, L.M., & Ton, J. (2016) The role of DNA (de)methylation in immune responsiveness of Arabidopsis. Plant J., 88, 361–374.

Lu, F., Cui, X., Zhang, S., Liu, C., & Cao, X. (2010) JMJ14 is an H3K4 demethylase regulating flowering time in Arabidopsis. Cell Res., 20, 387–390.

Luna, E., Bruce, T.J.A., Roberts, M.R., Flors, V., & Ton, J. (2012) Next-generation systemic acquired resistance. Plant Physiol., 158, 844–853.

Lyndsay, R.J., Holder, P.J., Talbot, N.J., & Gudelj, I. (2023) Metabolic efficiency reshapes the seminal relationship between pathogen growth rate and virulence. Ecol. Lett., doi: 10.1111/ele.14218.

Marazzi, I., Ho, J.S.Y., Kim, J., Manicassamy, B., Dewall, S., Albrecht, R.A., et al. 2012. Suppression of the antiviral response by an influenza histone mimic. Nature, 483, 428–433.

Martínez, F., Carrasco, J.L., Toft, C., Hillung, J., Giménez-Santamarina, S., Yenush, L., et al. 2023. A binary interaction map between turnip mosaic virus and *Arabidopsis thaliana* proteomes. *Commun*. Biol., 6, 28.

McCue, A.D., Nithikattu, S., Reeder, S.H., & Slotkin, R.K. (2012) Gene expression and stress response mediated by the epigenetic regulation of a transposable element small RNA. PLoS Genet., 8, e1002474.

McKenna, A., Hanna, M., Banks, E., Sivachenko, A., Cibulskis, K., Kernytsky, A., et al. 2010. The genome analysis toolkit: A MapReduce framework for analyzing next-generation DNA sequencing data. Genome Res., 20: 1297–1303.

Melero, I., González, R., & Elena, S.F. (2023) Host developmental stages shape the evolution of a plant RNA virus. Phil. Trans. R. Soc. B., 378, 20220005.

Mendizabal, I., Keller, T.E., Zeng, J., & Yi, S.V. (2014) Epigenetics and evolution. Integr. Compar. Biol., 54, 31–42.

Mongelli, V., Lequime, S., Kousathanas, A., Gausson, V., Blanc, H., Nigg, J., et al. (2022) Innate immunity pathways act synergistically to constrain RNA virus evolution in *Drosophila melanogaster*. *Nat*. Ecol. Evol., 6, 565–578.

Morel, J.B., & Dangl, J.L. (1997) The hypersenstivie response and the induction of cell death in plants. Cell Death Differ., 4, 671–683.

Mourrain, P., Béclin, C., Elmayan, T., Feuerbach, F., Godon, C., Morel, J.B., et al. (2000) Arabidopsis *SGS2* and *SGS3* genes are required for posttranscriptional gene silencing and natural virus resistance. Cell, 101, 533–542.

Moury, B., Morel, C., Johansen, E., Guilbaud, L., Souche, S., Ayme, V., et al. (2007) Mutations in *Potato virus Y* genome-linked protein determine virulence toward recessive resistances in *Capsicum annuum* and *Lycopersicon hirsutum*. Mol. Plant Microbe Interact., 17, 322–329.

Murata, T., Sugimoto, A., Inagaki, T., Yanagi, Y., Watanabe, T., Sato, Y., et al. (2021) Molecular basiss of Epstein-Barr virus latency establishment and lytic reactivation. Viruses, 13, 2344.

Murphy, J.F., Klein, P.G., Hunt, A.G., & Shaw, J.G. (1996) Replacement of the tyrosine residue that links a potyviral VPg to the viral RNA is lethal. Virology, 220, 535–538.

Navarro, R., Ambrós, S., Butković, A., Carrasco, J.L., González, R., Martínez, F., et al. (2022) Defects in plant immunity modulate the rates and patterns of RNA virus evolution. Virus Evol., 8, veac059.

Nei, M. (1982) Evolution of human races at the gene level. In: Bonné-Tamir B (ed.) Human Genetics, Part A: The Unfolding Genome. New York: Alan R. Liss. 167–181.

Nicaise, V., Gallois, J.L., Chafiai, F., Allen, L.M., Schurdi-Levraud, V., Browning, K.S., et al. (2007) Coordinated and selective recruitment of eIF4E and eIF4G factors for potyvirus infection in *Arabidopsis thaliana*. FEBS Lett., 581, 1041–1046.

Nuthikattu, S., McCue, A.D., Panda, K., Fultz, D., DeFraia, C., Thomas, E.N., et al. (2013) The initiation of epigenetic silencing of active transposable elements is triggered by RDR6 and 21-22 nucleotide small interfering RNAs. Plant Physiol., 162, 116–131.

Pagán, I., Alonso-Blanco, C., & García-Arenal, F. (2007) The relationship of within-host multiplication and virulence in a plant-virus system. PLoS ONE, 8, e786.

Pagán, I., Alonso-Blanco, C., & García-Arenal, F. (2008) Host responses in life-history traits and tolerance to virus infection in *Arabidopsis thaliana*. PLoS Pathog. 4, e1000124.

Pagán, I., Fraile, A., Fernández-Fueyo, E., Montes, N., Alonso-Blanco, C., & García-Arenal, F. (2010) *Arabidopsis thaliana* as a model for the study of plant-virus co-evolution. Phil. Trans. R. Soc. B, 365, 1938–1995.

Pettersen, E.F., Goddard, T.D., Huang, C.C., Meng, E.C., Couch, G.S., Croll, T.I., et al. (2021) UCSF ChimeraX: structure visualization for researchers, educators and developers. Prot. Sci., 30, 70–82.

Pontier, D., Picart, C., Roudier, F., García, D., Lahmy, S., Azevedo, J., et al. (2012) NERD, a plant-specific GW protein, defines an additional RNAi-dependent chromatin-based pathway in Arabidopsis. Mol. Cell, 48, 121–132.

Pontier, D., Yahubyan, G., Vega, D., Bulski, A., Saez-Vasquez, J., Hakimi, M.A., et al. (2005) Reinforcement of silencing at transposons and highly repeated sequences requires the concerted action of two distinct RNA polymerases IV in Arabidopsis. Genes Develop., 19, 2030–2010.

Rapp, R.A., & Wendel, J.E. (2005) Epigenetics and plant evolution. New Phytol., 168, 81–91.

Richard, C.L., Bossdorf, O., & Pigliucci, M. (2010) What role does heritable epigenetic variation play in phenotypic evolution? BioScience, 60, 232–237.

Rozas, J., Ferrer-Mata, A., Sánchez-del Barrio, J.C., Guirao-Rico, S., Librado, P., Ramos-Osins, S.E., et al. (2017) DnaSP v6: DNA sequence polymorphism analysis of large datasets. Mol. Biol. Evol., 34, 3299–3302.

Saze, H., Shiraihi, A., Miura, A., & Kakutani, T. (2008) Control of genic DNA methylation by a jmjC domain containing protein in *Arabidopsis thaliana*. Science, 319, 462–465.

Schmid-Hempel, P. (2013) Host-parasite genetics. In: Evolutionary Parasitology: The Integrated Study of Infections, Immunology, Ecology, and Genetics. Oxford: Oxford University Press. 259–272

Searle, I.R., Pontes, O, Melnyk, C.W., Smith, L.M., & Baulcombe, D.C. (2010) JMJ14, a JmjC domain protein, is required for RNA silencing and cell-to-cell movement of an RNA silencing signal in Arabidopsis. Genes Develop., 24, 986–991.

Shao, W., Kearney, M.F., Boltz, V.F., Spindler, J.E., Mellors, J.W., Maldarelli, F., et al. (2014) PAPNC, a novel method to calculate nucleotide diversity from large scale next generation sequencing data. J. Virol. Methods, 203, 73–80.

Singh, N., & Tscharke, D.C. (2020) Herpes simplex virus latency is noisier the closer we look. J. Virol. 94, e01701–19.

Simko, I., & Piepho, H. (2012) The area under the disease progress stairs: calculation, advantage, and application. Phytopathology, 102, 381–389.

Soosaar, J.L.M., Burch-Smith, T.M., & Dinesh-Kumar, S.P. (2005) Mechanisms of plant resistance to viruses. Nat. Rev. Microbiol., 3, 789–799.

Stajic, D., & Jansen, L.E.T. (2021) Empirical evidence for epigenetic inheritance driving evolutionary adaptation. Phil. Trans. R. Soc. B, 376, 20201421.

Stajic, D., Perfeito, L., & Jansen, L.E.T. (2019) Epigenetic gene silencing alters the mechanisms and rate of evolutionary adaptation. *Nat*. Ecol. Evol., 3, 4981–498.

Tajima, F. (1989) Statistical methods for testing the neutral mutation hypothesis by DNA polymorphism. Genetics, 123, 585–595.

Tamura, K., Stecher, G., & Kumar, S. (2021) Molecular Evolutionary Genetics Analysis version 11. Mol. Biol. Evol., 38, 3022–3027.

Traw, M.B., Kniskern, J.M., & Bergelson, J. (2007) SAR increases fitness of *Arabidopsis thaliana* in presence of natural bacterial pathogens. Evolution, 61, 2444–2449.

Voinnet, O. (2001) RNA silencing as a plant immune system against viruses. Trends Genet., 17, 449–459.

Wilm, A., Aw, P.P.K., Bertrand, D., Yeo, G.H.T., Ong, S.H., Wong, C.H., et al. (2012) LoFreq: A sequence-quality aware, ultra-sensitive variant caller for uncovering cell-population heterogeneity from high-throughput sequencing datasets. Nucleic Acids Res., 40, 11189–11201.

Wobbrock, J.O., Findlater, L., Gergle, D., & Higgins, J.J. (2011) The aligned rank transform for nonparametric factorial analysis using only ANOVA procedures. In Proceedings of the. ACM Conference on Human Factors in Computing Systems. New York: ACM Press, pp. 143–146.

Xie, Z., Johansen, L.K., Gustafson, A.M., Kasschau, K.D., Lellis, A.D., Zilberman, D., et al. (2004) Genetic and functional diversification of small RNA pathways in plants. PLoS Biol., 2, e104.

Yang. Z. (2006) Computational Molecular Evolution. UK: Oxford University Press, Oxford. pp. 262–264.

Yu, A., Lepère, G., Jay, F., Wang, J., Bapaume, L., Wang, Y., et al. (2013) Dynamics and biological relevance of DNA demethylation in Arabidopsis antibacterial defense. Proc. Natl. Acad. Sci. USA, 110, 2389–2394.

Zilberman, D., Cao, X., & Jacobsen, S.E. (2003) *ARGONAUTE4* control of locus-specific siRNA accumulation and DNA and histone methylation. Science, 299, 716–719.

Zou, J.M., & Zhang, Y. (2020) Plant immunity: danger perception and signaling. Cell, 181, 978–989.

