## Supplementary Information for "Phenotypic and genomic changes during *Turnip mosaic virus* adaptation to *Arabidopsis thaliana* mutants lacking epigenetic regulatory factors"

### Supplementary Material

**Figure S1**. Disease-related phenotypes measured for all 20 *A. thaliana* genotypes infected with TuMV, ordered according the four phenogroups defined in Table S1. (A) Incidence curves showing the frequency of infected plants along the 17 dpi. Infectivity values were estimated as the frequency of infection at 17 dpi; *AUDPS* values were estimated as the area under the corresponding incidence curves. (B) Symptoms development curves showing the evolution of disease severity during 14 dpi. Symptom severity was estimated as the mean values observed at 14 dpi; *AUSIPS* values are estimated as the area under the corresponding symptoms progression curves. (C) Increase in virus accumulation during the course of infections. Virus accumulation was evaluated by absolute RT-qPCR and expressed as fold-changes with respect to the corresponding values at 4 dpi. In all panels, dots represent the mean across plants and error bars represent ±1 SEM.

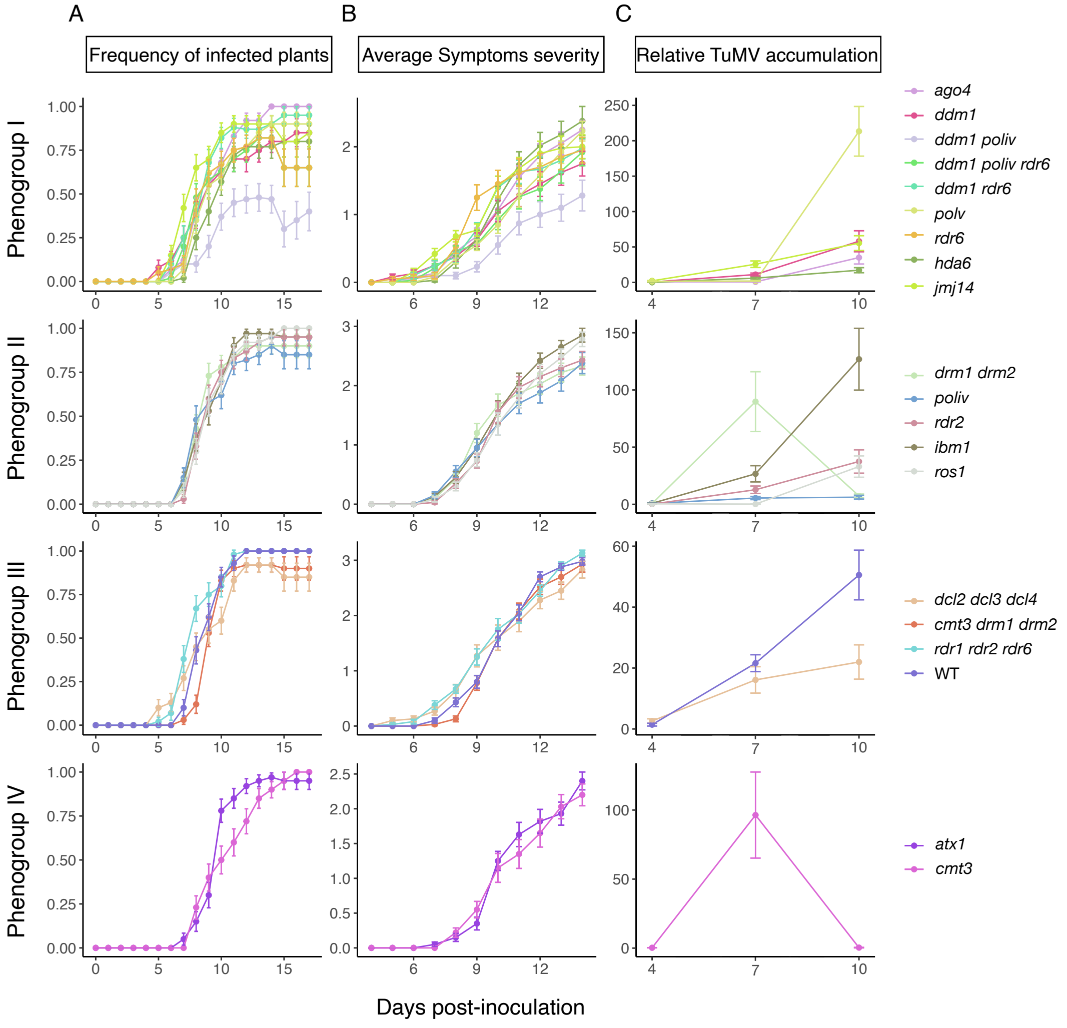

**Figure S2.** Schematic representation of the evolution experiment.

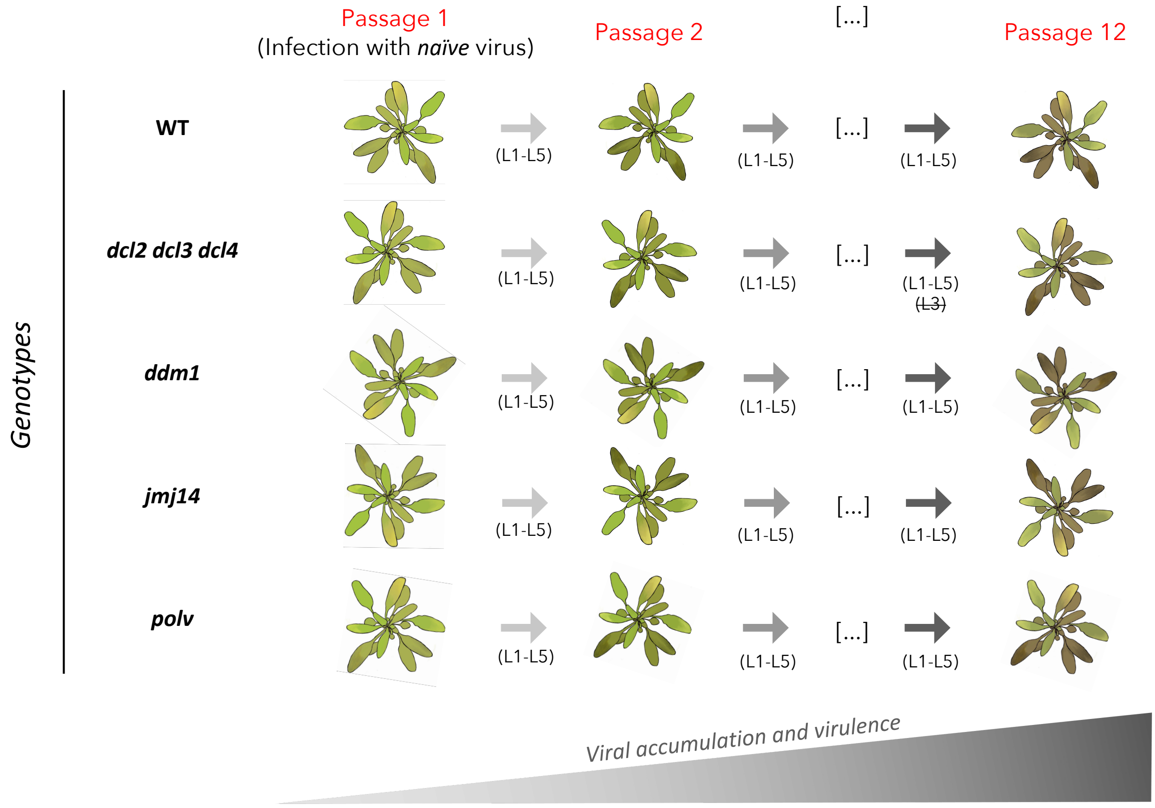

**Figure S3.** Evolution of the association between disease-related traits and log-viral load, measured as partial correlations controlling for plant genotype.

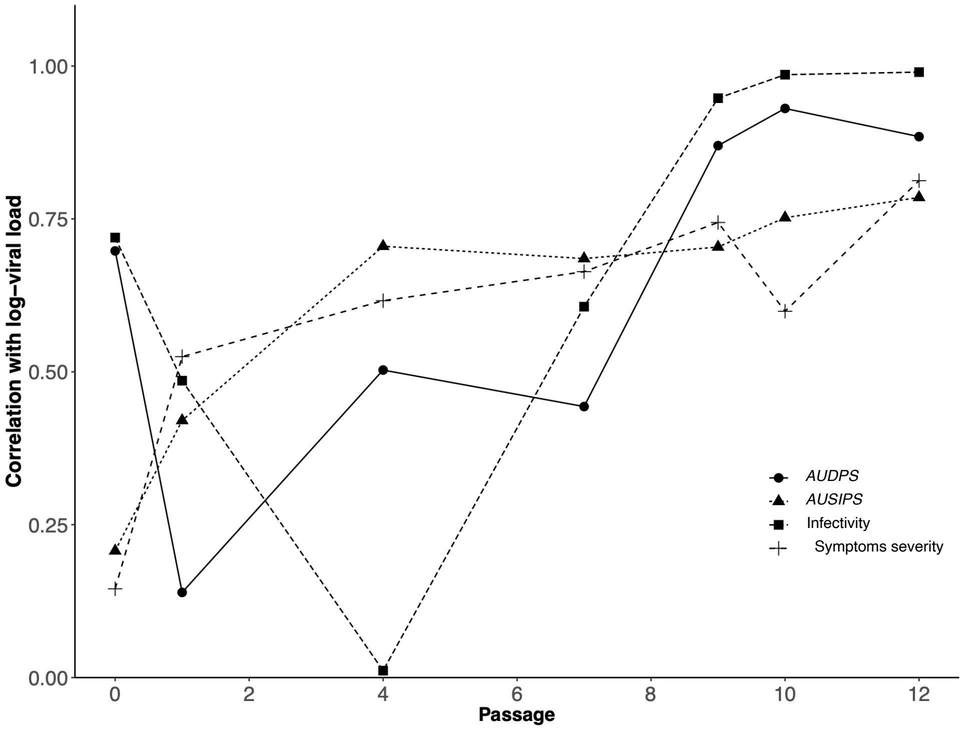

**Figure S4.** Distribution of variability, measured as Shannon's entropy, along TuMV genome at passages 1 and 12. Viral cistrons are indicated with different colors and replicates with different symbols (see legend).

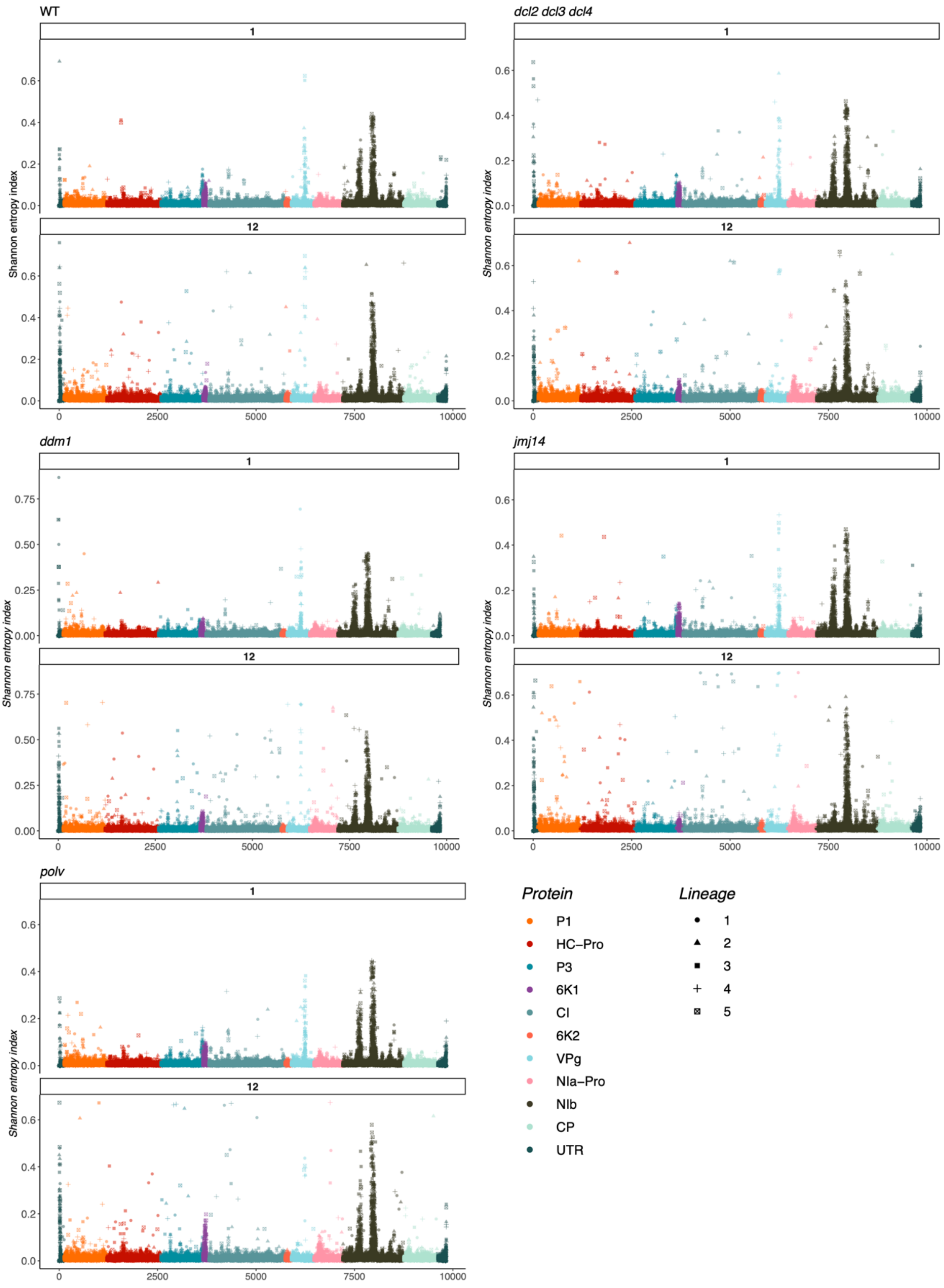

**Table S1**. Evaluation of the goodness-of-fit of models with an increasing number of clusters (phenogroups).

| Number of clusters | ***RSS***[1] | **–Log-likelihood** | ***BIC***[2] | **Akaike's weight** |
|---|---|---|---|---|
| 2 | 0.1272 | 18.5553 | 42.8913 | 0.0069 |
| 3 | 0.0934 | 14.2237 | 37.1185 | 0.1235 |
| 4 | 0.0831 | 11.1940 | 33.9496 | 0.6022 |
| 5 | 0.0372 | 11.8517 | 38.1553 | 0.0735 |
| 6 | 0.0332 | 10.2191 | 37.7803 | 0.0887 |
| 7 | 0.0316 | 8.8809 | 37.9944 | 0.0770 |
| 8 | 0.0166 | 9.2247 | 41.5723 | 0.0133 |
| 9 | 0.0157 | 8.3036 | 42.6206 | 0.0079 |
| 10 | 0.0157 | 7.4735 | 43.8507 | 0.0043 |

[1]Residual sum of squares

[2]In all cases, $n = 20$ genotypes.

**Table S2**. Results of the MANCOVA analysis fitting multivariate data to equation (1).

| **Effect** | $\boldsymbol{\Lambda}$[1] | **d.f.**[2] | $\boldsymbol{F}$ | $\boldsymbol{P}$ | $\boldsymbol{\eta_P^2}$ | $\mathbf{1-\boldsymbol{\beta}}$ |
|---|---|---|---|---|---|---|
| Intersection (*A*) | 0.9852 | 4, 272 | 4520.6449 | < 0.0001 | 0.9852 | 1 |
| Plant genotype (*G*) | 0.8826 | 16, 831.6 | 2.1682 | 0.0050 | 0.0307 | 0.9248 |
| Passage (*t*) | 0.4850 | 4, 272 | 72.2124 | < 0.0001 | 0.5150 | 1 |
| Passage by Plant genotype by ($t \times G$) | 0.7791 | 16, 831.6 | 4.4256 | < 0.0001 | 0.0605 | 0.9996 |
| Lineage within Plant genotype ($L(G)$) | 0.8823 | 80, 1075.4 | 0.4336 | 1 | 0.0308 | 0.7272 |
| Passage by Lineage within Plant genotype ($t \times L(G)$) | 0.4314 | 80, 1075.4 | 3.1933 | < 0.0001 | 0.1896 | 1 |

[1]Wilk's $\Lambda$.

[2]numerator, denominator

**Table S3**. Results of the ANCOVA analysis fitting log-viral load data to equation (1).

| Effect | $SS$[1] | d.f. | $F$ | $P$ | $\eta_P^2$ | $1-\beta$ |
|---|---|---|---|---|---|---|
| Intersection ($A$) | 2256.7749 | 1 | 229435.0165 | < 0.0001 | 1 | 1 |
| Plant genotype ($G$) | 0.6842 | 4 | 17.3899 | < 0.0001 | 0.7767 | 1 |
| Passage ($t$) | 9.9879 | 1 | 48.2857 | < 0.0001 | 0.2786 | 1 |
| Passage by Plant genotype by ($t \times G$) | 4.5256 | 4 | 5.4697 | 0.0004 | 0.1490 | 0.9720 |
| Lineage within Plant genotype ($L(G)$) | 0.1967 | 20 | 0.0476 | 1 | 0.0076 | 0.0675 |
| Passage by Lineage within Plant genotype ($t \times L(G)$) | 31.6424 | 20 | 7.6487 | < 0.0001 | 0.5503 | 1 |
| Error | 25.8562 | 125 | | | | |

[1]Type III sum of squares.

**Table S4**. Results of the MANOVA analysis fitting log-viral load data to equation (3).

| **Effect** | $\boldsymbol{\Lambda}$[1] | **d.f.**[2] | $\boldsymbol{F}$ | $\boldsymbol{P}$ | $\boldsymbol{\eta_P^2}$ | $\boldsymbol{1-\beta}$ |
|---|---|---|---|---|---|---|
| Intersection ($v$) | 0.0113 | 5, 16 | 2798.7670 | < 0.0001 | 0.9887 | 1 |
| Epigenetic pathway ($P$) | 0.1893 | 10, 32 | 4.1550 | 0.0010 | 0.5649 | 0.9883 |
| Plant genotype within pathway ($G(P)$) | 0.3869 | 10, 32 | 1.9443 | 0.0752 | 0.3780 | 0.7574 |

[1]Wilk's $\Lambda$.

[2]numerator, denominator.
